# A precision oncology-focused deep learning framework for personalized selection of cancer therapy

**DOI:** 10.1101/2024.12.12.628190

**Authors:** Casey Sederman, Chieh-Hsiang Yang, Emilio Cortes-Sanchez, Tony Di Sera, Xiaomeng Huang, Sandra D. Scherer, Ling Zhao, Zhengtao Chu, Eliza R. White, Aaron Atkinson, Jadon Wagstaff, Katherine E. Varley, Michael T. Lewis, Yi Qiao, Bryan E. Welm, Alana L. Welm, Gabor T. Marth

**Affiliations:** Eccles Institute of Human Genetics, University of Utah, Salt Lake City, Utah, USA; Huntsman Cancer Institute, University of Utah, Salt Lake City, Utah, USA; Department of Oncological Sciences, University of Utah, Salt Lake City, UT, USA; Departments of Molecular and Cellular Biology and Radiology. Lester and Sue Smith Breast Center. Dan L Duncan Comprehensive Cancer Center. Baylor College of Medicine, Houston, Texas, USA; Department of Biomedical Informatics, School of Medicine, University of Utah, Salt Lake City, UT, USA; Department of Surgery, University of Utah, Salt Lake City, UT, USA

## Abstract

Precision oncology matches tumors to targeted therapies based on the presence of actionable molecular alterations. However, most tumors lack actionable alterations, restricting treatment options to cytotoxic chemotherapies for which few data-driven prioritization strategies currently exist. Here, we report an integrated computational/experimental treatment selection approach applicable for both chemotherapies and targeted agents irrespective of actionable alterations. We generated functional drug response data on a large collection of patient-derived tumor models and used it to train ScreenDL, a novel deep learning-based cancer drug response prediction model. ScreenDL leverages the combination of tumor omic and functional drug screening data to predict the most efficacious treatments. We show that ScreenDL accurately predicts response to drugs with diverse mechanisms, outperforming existing methods and approved biomarkers. In our preclinical study, this approach achieved superior clinical benefit and objective response rates in breast cancer patient-derived xenografts, suggesting that testing ScreenDL in clinical trials may be warranted.

## Main

Widespread adoption of clinical genetic testing, married with an expanding repertoire of anticancer therapies, has significantly expanded the reach of precision cancer treatments.^1^ However, despite several successful applications of genome-guided therapy, such as the approval of EGFR inhibitors in EGFR-mutant non-small cell lung cancer and PARP inhibitors in specific tumor types harboring BRCA1/2 mutations, a lack of rational guidance for treatment selection remains a critical barrier to precision oncology practice. Most drugs lack approved pretreatment biomarkers,^1^ and most tumors lack clinically actionable molecular alterations, limiting biomarker-informed treatment selection to a minority of patients.^1–4^ Indeed, the pioneering NCI-MATCH trial of >5,900 patients with diverse malignancies identified actionable alterations in just 38% of individuals, with <18% ultimately assigned to treatment arms based on genomic indicators.^5^ Further, response to single-gene biomarker-informed treatments is highly variable, even among patients harboring theoretically targetable alterations.^6–8^ These challenges are particularly acute in metastatic tumors with their vast inter-tumoral heterogeneity and diverse resistance mechanisms.^9^ While recent metastatic cancer trials identified actionable alterations in 40-46% of patients, no clinical benefit was observed when matching patients to genome-guided therapies.^10,11^ Given these limitations, chemotherapy remains a mainstay of cancer treatment.^3,4^ However, no data-driven approaches exist to guide selection across the full spectrum of relevant chemotherapeutic agents.^12^ Taken together, these considerations suggest that expanding the currently limited scope of precision oncology practice requires: (1) extending precision treatment selection to drugs, including chemotherapies, lacking validated predictive biomarkers^1,13^ and (2) refining the prediction of targeted agent sensitivity beyond the presence or absence of single-gene molecular alterations.

To this end, recent advances in deep learning (DL)-based cancer drug response prediction (CDRP) offer an alternative to today’s biomarker-based treatment selection strategies.^14^ These approaches leverage large-scale pharmaco-omic screens in cancer cell lines to train data-driven models that can be applied to predict treatment response in patient tumors. Critically, DL approaches incorporate high-dimensional omic features (e.g., uncurated lists of somatic mutations or gene expression values) without the explicit specification of known prognostic biomarkers, facilitating response prediction for drugs lacking validated biomarkers and for tumors lacking actionable alterations.^14,15^ However, while existing models perform well in standard cell line benchmarks, performance declines significantly under precision oncology-relevant testing conditions. Indeed, a recent study found that existing DL models suffer from an overreliance on drug-level features, ultimately preventing the generation of personalized predictions based on a tumor’s omic characteristics.^16^ Further, compared to human tumors, cell lines exhibit differences in key biological variables associated with drug response, including differentiation, proliferation, and drug metabolism.^17,18^ As a consequence, pharmaco-omic associations learned in cell lines do not necessarily translate to patients in a straightforward manner, limiting the clinical utility of models trained in cell lines alone.

To better model human tumors, we and others have advanced a series of patient-derived models of cancer (PDMCs) maintaining strong biological fidelity and concordant drug responses with the originating patient’s tumor.^19–21^ In particular, patient-derived organoids (PDOs)^22^ and other short-term *ex vivo* culture systems^23,24^ have received significant attention, serving as intermediate models between *in vitro* cell lines and *in vivo* xenografts while enabling functional drug screening within a clinically relevant timeframe^21,25^. Beyond providing a more clinically relevant source of pharmaco-omic data for CDRP models, functional testing in PDMCs has proven a powerful standalone tool for treatment selection in multiple cancer types.^22,26^ Further, PDMC-based functional testing has identified targeted agent sensitivities in tumors lacking the associated therapeutic biomarkers^27,28^ and revealed distinct drug response profiles in tumors harboring similar driver mutations.^25^ The ability of functional testing to augment tumor omic information has motivated recent trials pairing tumor omic characterization with PDMC-based functional testing to provide complementary views of drug sensitivity that can be integrated to inform treatment decisions.^29^ However, existing integration strategies rely on *post hoc* correlative analyses, necessitating agreement between omic and functional findings. While concordant results can increase confidence when selecting treatments, a lack of established practices to resolve contradictory findings limits the synergy between omic and functional modalities.

To address these gaps, here we report a unified computational/experimental approach to treatment selection, combining a clinically oriented functional precision oncology (FPO) pipeline with ScreenDL, a novel DL-based CDRP framework engineered to exploit the patient-relevant data generated by this experimental protocol. During training, ScreenDL progressively incorporates data of increasing patient relevance to generalize pharmaco-omic associations previously learned in cell lines for response prediction in patients. As new patients enter our FPO pipeline, parallel tumor omic profiling and functional drug screening in matched organoids enables patient-specific fine-tuning with our ScreenAhead module, integrating a tumor’s omic and functional characteristics with prior knowledge learned from training examples to generate personalized response predictions. ScreenDL outperforms existing DL models across a panel of clinically relevant benchmarks in cell lines and PDMCs. Further, personalization with ScreenAhead dramatically improves response prediction, highlighting the power of our combined computational/experimental strategy. Mirroring clinical application in patients and matched PDOs, we applied our end-to-end computational/experimental treatment selection framework to high-risk/metastatic patient-derived xenograft (PDX) models with functional profiling in matched PDX-derived organoids (PDXOs). Our promising results in this preclinical validation justify further evaluation in future clinical trials to aid precision treatment selection in the clinic.

## Results

### Design and training of a deep learning model optimized for precision oncology applications

Cancer drug response involves the interplay of complex biological and chemical factors.^30^ To model these interactions, ScreenDL learns a function *R = f(D, T)*, mapping a drug’s chemical structure *D* and a tumor’s transcriptomic profile *T* to a predicted response *R* (Fig. 1a). In this schema, drugs are encoded as Morgan fingerprints,^31^ a canonical vector representation of chemical structure, and tumors are represented as vectors encoding the *z*-score normalized expression of 4,364 genes from the Molecular Signatures Database (MSigDB) hallmark gene set collection^32^ (Fig. 1a). Here, the use of hallmark genes corresponding to well-defined biological processes^32^ allows ScreenDL to link cancer-relevant cellular phenotypes with drug response characteristics while minimizing reliance on individual genomic biomarkers for prediction. These tumor and drug encodings are passed to parallel feature extraction subnetworks dedicated to learning rich embeddings of a tumor’s transcriptomic profile and a drug’s chemical structure, respectively. The resulting embeddings are then concatenated and fed to a shared response prediction subnetwork which models interactions between biological and chemical features to predict therapy response. Drug response is quantified as *z*-score ln(IC50), termed *Z_D_*, corresponding to the *z*-score normalized, log transformed half-maximal inhibitory concentration of a given tumor-drug pair. *Z*-score normalization was performed independently for each drug across tumor samples, removing drug-specific biases in standard dose-response metrics^16,33^ and promoting transcriptomic signals associated with differential response - that is, the response of a given tumor to a given drug relative to other tumors. This formulation allows ScreenDL to identify therapies that elicit exceptional responses in the tumor of interest.

**Fig. 1:**
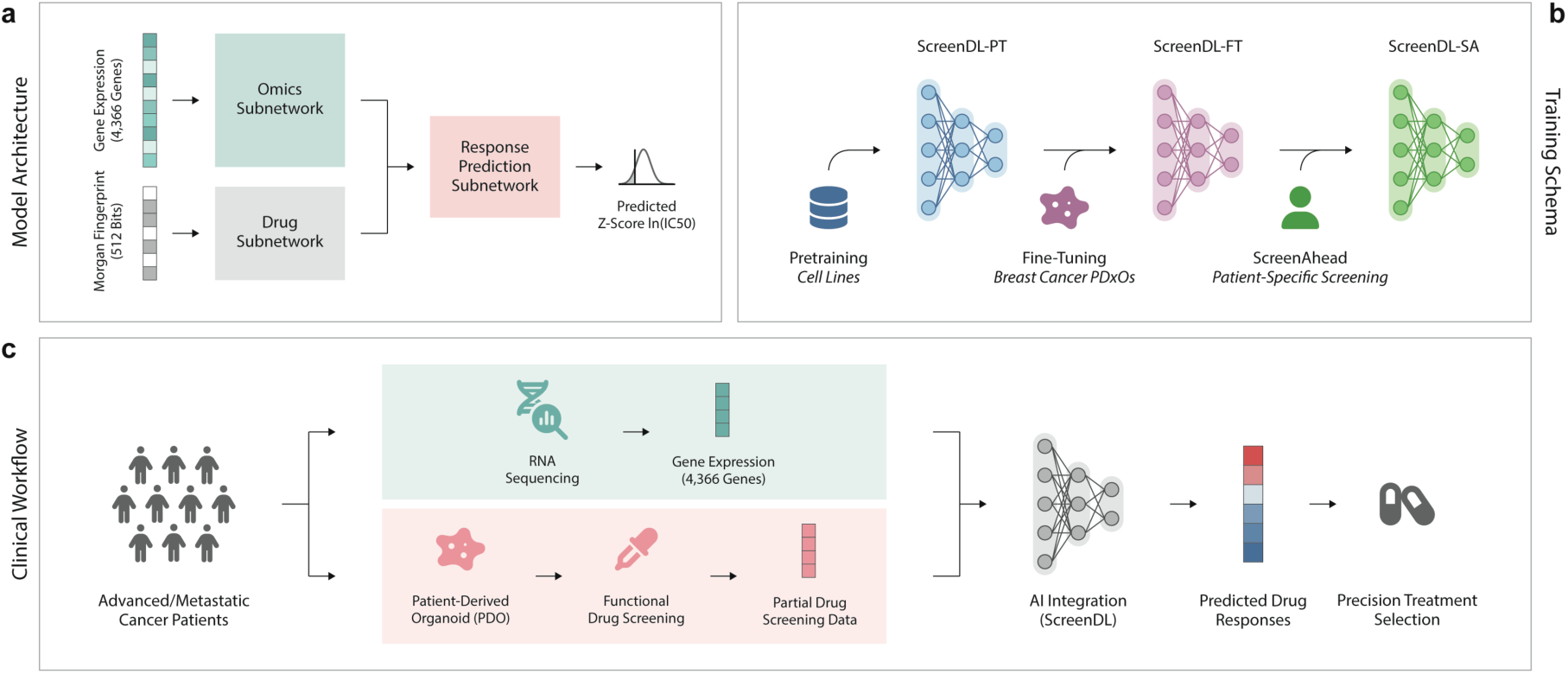
Deep learning-based integration of tumor omic and functional data for precision treatment selection with ScreenDL. **a.** Model Architecture: ScreenDL takes a tumor’s transcriptomic profile and a drug’s chemical structure as input and predicts z-score ln(IC50) values. **b.** Data Sources and Training Schema: During initial general-purpose pretraining, ScreenDL extracts generalizable associations between tumor omics and drug response from large-scale cell line pharmaco-omic databases (blue). During subsequent domain-specific fine-tuning, ScreenDL leverages pharmaco-omic data from breast cancer PDXOs to adapt to a more clinically relevant response prediction context (purple). Finally, patient-specific fine-tuning with ScreenAhead generates a personalized response prediction model optimized for the N-of-1 precision oncology context (green). **c.** Clinical Workflow: When a new patient enters the functional precision oncology pipeline, a biopsy is taken, and RNA sequencing is performed. In parallel, a patient-derived organoid model is established, and functional drug screening is performed. The resulting multimodal data is integrated with prior knowledge through patient-specific fine-tuning with ScreenAhead. Predicted drug responses are then used to select an optimal treatment.

The training of ScreenDL proceeds in three phases, each designed to maximize available pharmaco-omic data and provide accurate response predictions for never-before-seen tumor samples (Fig. 1b). During an initial ***pretraining*** phase, ScreenDL leverages large-scale pharmaco-omic screens performed in cell lines to extract generalizable associations between tumor omics and drug response. To establish a comprehensive pharmaco-omic dataset for pretraining, we harmonized drug screening data from the Genomics of Drug Sensitivity in Cancer database^34^ with matched transcriptomic profiles from Cell Model Passports^35^. In total, this harmonized dataset consists of 278,033 cell line-drug pairs spanning 799 pan-cancer cell lines and 409 anticancer therapies. Following general-purpose pretraining in cell lines, context-specific pharmaco-omic data derived from more clinically relevant PDXOs was integrated through ***domain-specific fine-tuning***, adapting the pretrained ScreenDL model to a more clinically relevant response prediction context. This fine-tuning schema allows ScreenDL to capitalize on less-abundant, but highly patient-relevant pharmaco-omic data from PDMCs while retaining the generalizable knowledge learned during pretraining. In the last phase, personalization was achieved through ***patient-specific fine-tuning*** with our ScreenAhead module. This step integrates patient-level transcriptomic features with limited functional drug screening in PDMCs derived from the patient to generate a personalized response prediction model (Fig. 1c). While existing CDRP frameworks rely solely on tumor omic features for inference^14^, our ScreenAhead strategy incorporates dual omic and functional characteristics (Fig. 1c), allowing ScreenDL to exploit partial functional drug screening to augment tumor omic inputs and enhance predictions for a greater number of drugs.

### ScreenDL provides accurate response predictions in never-before-seen cancer cell lines

As an initial assessment, we measured the prediction accuracy of our pre-trained ScreenDL model (ScreenDL-PT) in never-before-seen cell lines using the Pearson correlation coefficient (PCC) between observed and predicted *Z_D_* values (see Methods). To mitigate the propensity of global metrics to overstate the accuracy of CDRP models^16^, performance was assessed independently for each drug. Across all tested drugs, ScreenDL-PT achieved a median PCC of 0.47, a significant improvement compared to 0.43 for DeepCDR^36^ (p = 2.84 x 10^-63^, Wilcoxon signed-rank test), the best of three existing DL models: DeepCDR, DualGCN^37^, and HiDRA^38^ (Fig. 2a). Further, ScreenDL-PT outperformed all tested models regardless of treatment type, achieving median PCCs of 0.56 and 0.52 across chemotherapies and targeted agents, respectively (Fig. 2a). To glean additional insight, we further stratified performance, grouping drugs by biological mechanism and cell lines by parent tissue type. ScreenDL-PT outperformed all tested models across the breadth of drug mechanisms, with median PCCs ranging from 0.41 for drugs targeting receptor tyrosine kinase signaling to 0.59 for drugs targeting the ERK/MAPK axis (Fig. 2b). Further, ScreenDL-PT outperformed the next-best model for 63% of tissue types compared to just 19% for HiDRA, the next-best model (Fig. 2c).

**Fig. 2:**
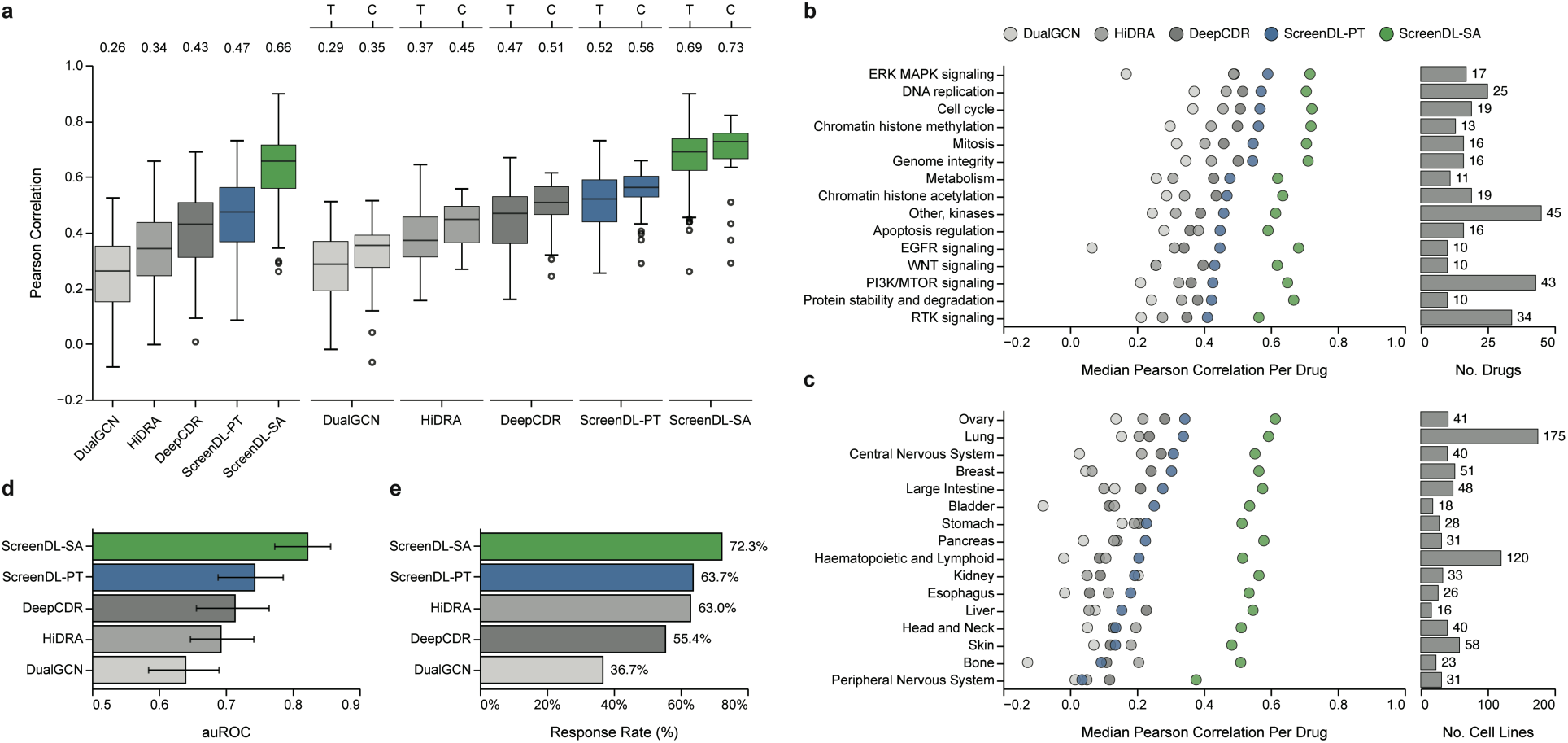
ScreenDL enables accurate response prediction in never-before-seen cell lines. **a.** Comparison of the drug level-prediction accuracies achieved by ScreenDL-PT and ScreenDL-SA with those of three existing DL-based CDRP models. Box plots represent the distribution of Pearson correlations between observed and predicted response per drug. Median values for each model are denoted in the upper margin. Drug-level performance is stratified by drug type for a subset of drugs (n = 213) which were manually annotated as either: chemotherapies (C), n = 34; or targeted agents (T), n = 179. **b.** Performance stratified according to drug biological mechanisms. Points represent the median Pearson correlation coefficient per drug. Bars indicate the number of training drugs per biological mechanism. Only mechanisms with at least 10 drugs are shown. **c.** Drug-level performance stratified according to cell line tissue types. Points represent the median Pearson correlation per drug amongst cell lines from the corresponding tissue. Bars indicate the number of training cell lines per tissue. Only tissues with at least 10 cell lines are shown. **d.** Median area under the receiver operating characteristic curve (auROC) across drugs for ScreenDL-PT, ScreenDL-SA, and three existing DL models. For each drug, sensitive tumors were defined as cell lines with an observed response falling below the 30th percentile of observed *Z_D_* values. Error bars denote interquartile ranges. **e.** Cell line response rates to drugs selected by different DL models.

To quantify ScreenDL’s ability to stratify sensitive and resistant tumors, we binarized observed *Z_D_* values and computed area under the receiver operating characteristic curve (auROC) independently for each drug. ScreenDL-PT achieved a median auROC of 0.74 compared to 0.71 for DeepCDR, the next best model (Fig. 2d). We next applied ScreenDL-PT to select optimal precision treatments for each cell line and quantified response rate (RR). Here, RR was defined as the fraction of cell lines for which response fell below the 30th percentile of observed *Z_D_* values for the selected therapy, a threshold predictive of partial or complete response in patients^23^. ScreenDL-PT achieved a RR of 64% amongst selected therapies, a slight improvement compared to HiDRA (RR = 63%) and the highest of all tested models (Fig. 2e). Closer inspection of the therapies selected by ScreenDL-PT revealed 171 anticancer agents targeting over 20 distinct cancer-associated biological pathways, illustrating ScreenDL’s ability to match cell lines with treatments tailored to their individual transcriptomic characteristics.

### Generation of an expansive high-risk/metastatic breast cancer PDMC pharmaco-omic resource

Despite these promising results in cell lines, ScreenDL-PT achieved only a modest median PCC of 0.27 across drugs when applied to our previously reported breast cancer PDXO collection^21^ (n = 16 PDXOs with pharmaco-omic data; Extended Data Fig. 1). While ScreenDL still outperformed DeepCDR (median PCC = 0.17) and HiDRA (median PCC = 0.15) in this PDXO cohort, the observed performance loss across all tested models underscores the limitations of cell lines when training DL models for response prediction in patient tumors. To provide a more extensive pharmaco-omic dataset to fine-tune our models, we generated an expanded living biobank of stable organoid lines (n = 99 PDXO + 1 PDO) from high-risk/metastatic breast tumors following our published protocols.^21,39^ Comprehensive descriptions of these PDXO lines and clinical annotations for the originating tumor samples are provided in Supplementary Table S1, and omic profiles of the originating patient tumor (when available), the PDX, and the matching PDXO lines were imported into cBioPortal.^40–42^ Each of the organoid lines was functionally characterized for response to a subset of >100 anticancer therapies centering on treatments approved for breast cancer and drugs available through the National Cancer Institute’s Cancer Therapy Evaluation Program. A summary of drugs tested in each PDXO line is provided in Supplementary Table S2. This effort yielded an extensive pharmaco-omic dataset including nearly 5,000 drug response observations spanning 100 PDXO/PDO lines and >100 anticancer therapies (Fig. 3a; Supplementary Fig. 1a-d); Supplementary Table S2).

**Fig. 3:**
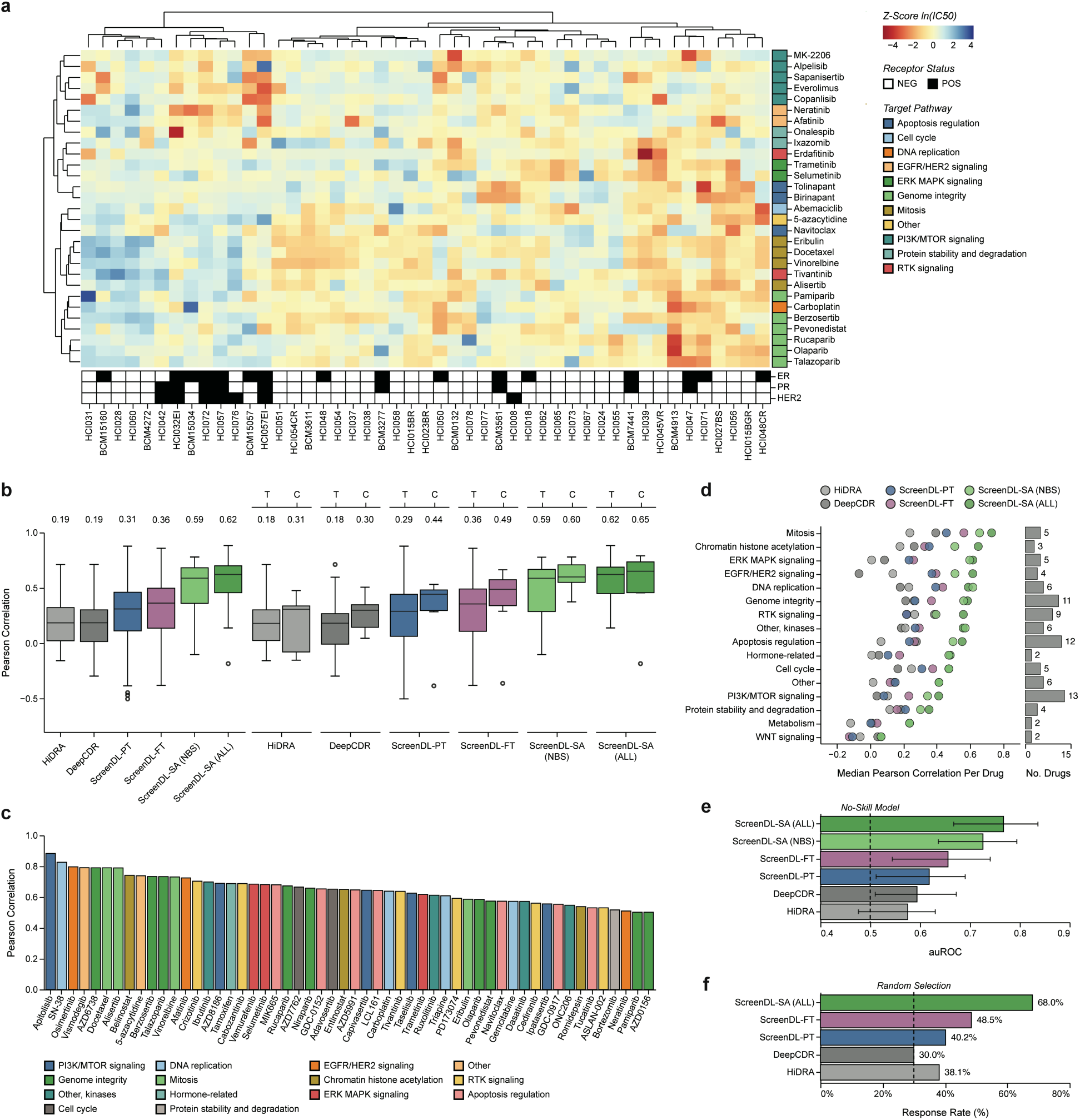
ScreenDL achieves accurate response prediction in high-risk/metastatic breast cancer PDXO models. **a.** Unsupervised clustering of 29 drugs carried forward for screening in at least 70% of PDXO lines. Only the 48 PDXO lines in which all 29 drugs were screened are included. Color indicates *z*-score normalized ln(IC50) values (darker red indicates cytotoxicity and darker blue indicates growth). Row and column annotations denote the biological mechanism of screened compounds and the hormone receptor status of each PDXO line, respectively. **b.** Performance of each ScreenDL variant in our expanded breast cancer PDXO cohort for the subset of 61 drugs included in cell line pretraining compared with that of two existing DL-based CDRP models. Box plots represent the distribution of Pearson correlations between observed and predicted response per drug. Median values for each model are denoted in the upper margin. Drug-level performance is stratified by drug type for a subset of drugs which were manually annotated as either: chemotherapies (C), n = 10; or targeted agents (T), n = 51. **c.** High-confidence drugs (PCC > 0.5) for ScreenDL-SA (ALL) colored by drug biological mechanisms. **d.** Drug-level performance stratified according to the biological mechanism of the corresponding therapy. Points represent the median Pearson correlation per drug. Bars indicate the number of training drugs per target pathway. Only biological mechanisms with at least two drugs are shown. **e.** Median area under the receiver operating characteristic curve (auROC) across drugs for ScreenDL-PT, ScreenDL-SA, and two existing DL models. For each drug, sensitive tumors were defined as PDXOs with an observed response falling below the 30th percentile of observed *Z_D_* values for a given drug. Error bars denote interquartile ranges. **f.** PDXO response rates for drugs selected by different DL models.

Unsupervised clustering of PDXO response profiles for the 29 drugs that showed promising activity and were therefore screened in at least 70% of PDXO lines revealed similar patterns of response to groups of drugs sharing common biological mechanisms, including microtubule inhibitors and drugs targeting PI3K/MTOR signaling (Fig. 3a). Similarly, response profiles for agents with the same protein targets were highly correlated, including responses to the PARP inhibitors olaparib and talazoparib (PCC = 0.72, p = 2.14 x 10^-15^) and the SMAC mimetics birinapant and tolinapant (PCC = 0.86, p = 1.06 x 10^-14^). PDXO responses also recapitulated established clinical biomarkers, with drugs targeting PI3K/MTOR signaling showing increased activity in estrogen receptor-positive (ER+) lines (p = 0.0014, Mann-Whitney U test) and enhanced sensitivity to platinum-based chemotherapies in triple-negative breast cancer lines (p = 0.009, Mann-Whitney U test).^43^ Finally, validation of 98 PDXO responses in the originating PDX samples revealed that PDXO screening was strongly predictive of *in vivo* treatment response (Extended Data Fig. 2), corroborating our earlier conclusions^21^ and illustrating the ability of this organoid platform to faithfully recapitulate *in vivo* drug response dynamics.

### Domain-specific fine-tuning improves response prediction in breast cancer PDMCs

We next investigated whether fine-tuning ScreenDL using this expanded PDXO pharmaco-omic dataset would improve model performance. To establish a baseline, we first assessed the prediction accuracy of ScreenDL-PT in these 100 PDXOs using the subset of 61 drugs also screened in cell lines. ScreenDL-PT achieved a median PCC across drugs of 0.31, outperforming both DeepCDR (median PCC = 0.19) and HiDRA (median PCC = 0.19) (Fig. 3b). Notably, while performance on drugs not included in cell line pretraining was comparatively poor (median PCC = 0.18; Extended Data Fig. 3), ScreenDL-PT still provides reliable predictions for a subset of such drugs, including the BRAF inhibitor vemurafenib (PCC = 0.65) and the microtubule inhibitor eribulin (PCC = 0.33). We then fine-tuned ScreenDL-PT using our full breast cancer PDXO dataset and assessed performance under leave-one-out cross-validation (see Methods). The fine-tuned model (ScreenDL-FT) displayed a significant increase in accuracy (PCC = 0.36; p = 0.003, Wilcoxon signed-rank test; Fig. 3b), with a mean improvement per drug of 0.06. Marked improvement was also observed for drugs not included in cell line pretraining (mean improvement per drug = 0.05; Extended Data Fig. 3). Further, fine-tuning improved performance for both targeted agents and chemotherapies (Fig. 3b) and for the vast majority of drug biological mechanisms (75%; Fig. 3f).

### Personalization with ScreenAhead significantly increases predictive power and improves treatment selection

To capitalize on our ability to generate matched omic and functional data for new patients through our FPO pipeline^21,44^, we tested whether tumor-specific fine-tuning (ScreenAhead) using a tumor’s transcriptomic profile and partial drug screening data could further improve performance. Here, we reasoned that personalization using a small, pre-screened drug panel would improve predictions across the broader space of unscreened therapies. As an initial test, we performed ScreenAhead tumor-specific fine-tuning for each cell line in our pharmaco-omic dataset and quantified prediction accuracy in unseen cell line-drug pairs. Specifically, for each cell line, a subset of 20 drugs was selected for ScreenAhead (see Methods). We then fine-tuned ScreenDL-PT using the cell line’s transcriptomic profile and its responses to these 20 drugs. ScreenAhead dramatically improved performance regardless of drug mechanism or tissue type (Fig. 2b,c), with ScreenDL-SA achieving a median PCC per drug of 0.66 compared to 0.47 for ScreenDL-PT (p = 1.15 x 10^-68^, Wilcoxon signed-rank test; Fig. 2a). Given potential biological and technical restrictions on the scope of pre-treatment drug screening, we explored the efficacy of our ScreenAhead approach while incrementally expanding the number of pre-screened drugs for each cell line. We observed significant improvements in drug-level performance with as few as 5 pre-screened drugs, highlighting ScreenAhead’s ability to exploit a minimal screening panel to improve predictions for unscreened agents (Extended Data Fig. 4a, Supplementary Text 1).

We next assessed ScreenAhead in our breast cancer PDXO cohort. Specifically, following domain-specific fine-tuning, ScreenAhead tumor-specific fine-tuning was performed for each PDXO using 12 drug responses (a feasible number to screen on PDOs in a trial setting) and the PDXO’s transcriptomic profile. ScreenDL-SA showcased superior prediction accuracy compared to ScreenDL-FT, achieving a median PCC per drug of 0.62 when considering all drug responses (ScreenDL-SA ALL; Fig. 3b) and 0.59 when considering only never-before-seen (NBS) PDXO-drug pairs, i.e., those PDXO-drug pairs not included in ScreenAhead (ScreenDL-SA NBS; Fig. 3b). Interrogation of drug-level performance revealed a subpopulation of drugs with exceptionally high prediction accuracy (57% of drugs with PCC > 0.50), including both chemotherapies and targeted agents with diverse mechanisms of action (Fig. 3c). Further, when restricted to PDXO-drug pairs not included in ScreenAhead, ScreenDL-SA still provides high-confidence predictions for 43% of drugs. Finally, ScreenDL-SA achieved a median auROC across drugs of 0.77 (Fig. 3e) and a 68% response rate (Fig. 3f), outperforming all competing models in both metrics. Together with ScreenDL-SA’s superior performance regardless of drug biological mechanism (Fig. 3d), these findings demonstrate the broad applicability of ScreenDL for response prediction in breast cancer patients.

### ScreenAhead harnesses two distinct types of information encoded in partial screening data

Motivated by the superior performance of ScreenDL-SA, we sought to understand how specifically ScreenAhead exploits partial pre-screening data to improve predictions. Given reports citing strong correlations between a tumor’s global drug sensitivity (GDS), defined as the tumor’s average response across drugs, and response to individual agents,^45,46^ we explored whether ScreenAhead leverages knowledge of GDS extracted from partial screening data to calibrate predictions for a given tumor. In both cell lines and PDXOs, GDS was strongly correlated with response to individual treatments (Extended Data Fig. 5a,b). However, tumor-specific fine-tuning using a cell line’s mean-filled *Z_D_* responses revealed that, while incorporating knowledge of GDS improved performance, GDS alone did not account for the full performance gain seen when using a cell line’s true *Z_D_* values in ScreenAhead (Extended Data Fig. 5c). Further, we found that ScreenAhead improved predictions for tumor-drug pairs for which GDS was not highly predictive, suggesting the ScreenAhead exploits additional information encoded in partial pre-screening data beyond a tumor’s GDS (Extended Data Fig. 5d,e; Supplementary Text 2).

An inspection of ScreenDL’s drug subnetwork embeddings revealed stratification according to common biological mechanisms (Extended Data Fig. 6d,e). Moreover, ScreenDL-SA’s performance gains were strongly correlated with the functional similarity of a given drug to drugs in a cell line’s pre-screening drug set (Extended Data Fig. 6a,b; Supplementary Text 3). These findings suggested that ScreenAhead might improve predictions for unscreened drugs by exploiting learned drug-drug functional similarities to borrow information from functionally related therapies included in pre-screening. To test this hypothesis, we performed a focused evaluation of ScreenDL-SA on 20 drugs when including an increasing number of functionally related therapies in ScreenAhead (see Methods). We observed a significant increase in drug-level performance when at least one functionally related therapy was included (p = 1.91 x 10^-6^, Wilcoxon signed-rank test; Extended Data Fig. 6c). Further improvement was observed with the addition of each subsequent functionally related therapy. These insights motivated a comparative analysis of ScreenAhead drug selection methods, revealing that informed drug selection using principal feature analysis (PFA)^47^ significantly outperformed both random drug selection and several alternative drug selection strategies (Extended Data Fig. 4b-d; Supplementary Text 1). Relative to random selection, the superior performance of PFA in this analysis supports the positive transfer of information across functionality-related therapies during ScreenAhead.

### ScreenDL outperforms single-gene biomarkers for targeted agent selection in cell lines and PDXOs

To date, precision oncology has focused on the development of genome-guided therapies targeting genetic alterations seen repeatedly in tumors.^48^ However, the presence of these genetic markers does not guarantee sensitivity to the corresponding targeted agents.^11^ To determine whether ScreenDL could improve patient stratification relative to single- or multi-gene biomarkers alone, we compared the performance of ScreenDL to biomarker-only predictive models. In cell lines, *BRAF* mutation status was strongly predictive of response to dabrafenib, a targeted *BRAF* inhibitor approved for specific cancers harboring oncogenic *BRAF* mutations (Fig 4a,b). However, a number of dabrafenib-sensitive cell lines lacked *BRAF* mutations and would not be selected for dabrafenib treatment based on mutational status alone. In comparison, ScreenDL-PT enables stratification of cell lines within genomically defined subgroups (Fig 4c). Further, ScreenAhead with just 20 drugs (excluding dabrafenib) dramatically improved performance, both overall (PCC = 0.72) and within genomic subgroups (PCC = 0.74, *BRAF* mutant cell lines; PCC = 0.63, *BRAF* wild-type cell lines; Fig 4d). We next considered capivasertib, an AKT inhibitor approved in combination with fulvestrant for hormone receptor positive, HER2 negative metastatic breast tumors harboring *PIK3CA, AKT1*, or *PTEN* mutations. While driver mutations in *PIK3CA, AKT1* or *PTEN* were associated with increased capivasertib sensitivity in cell lines (Fig. 4e), this association translated to comparatively poor performance in a biomarker-only model (PCC = 0.25; Fig. 4f). In comparison, ScreenDL-PT achieved a PCC of 0.50 while again enabling stratification within genomic subgroups, particularly for cell lines lacking the qualifying driver mutations where ScreenDL-PT recovered a subset (n = 31) of exceptionally capivasertib-sensitive lines (observed *Z_D_* below the 10th percentile) that lacked *PIK3CA, AKT1*, or *PTEN* driver mutations (Fig. 4g).

**Fig. 4:**
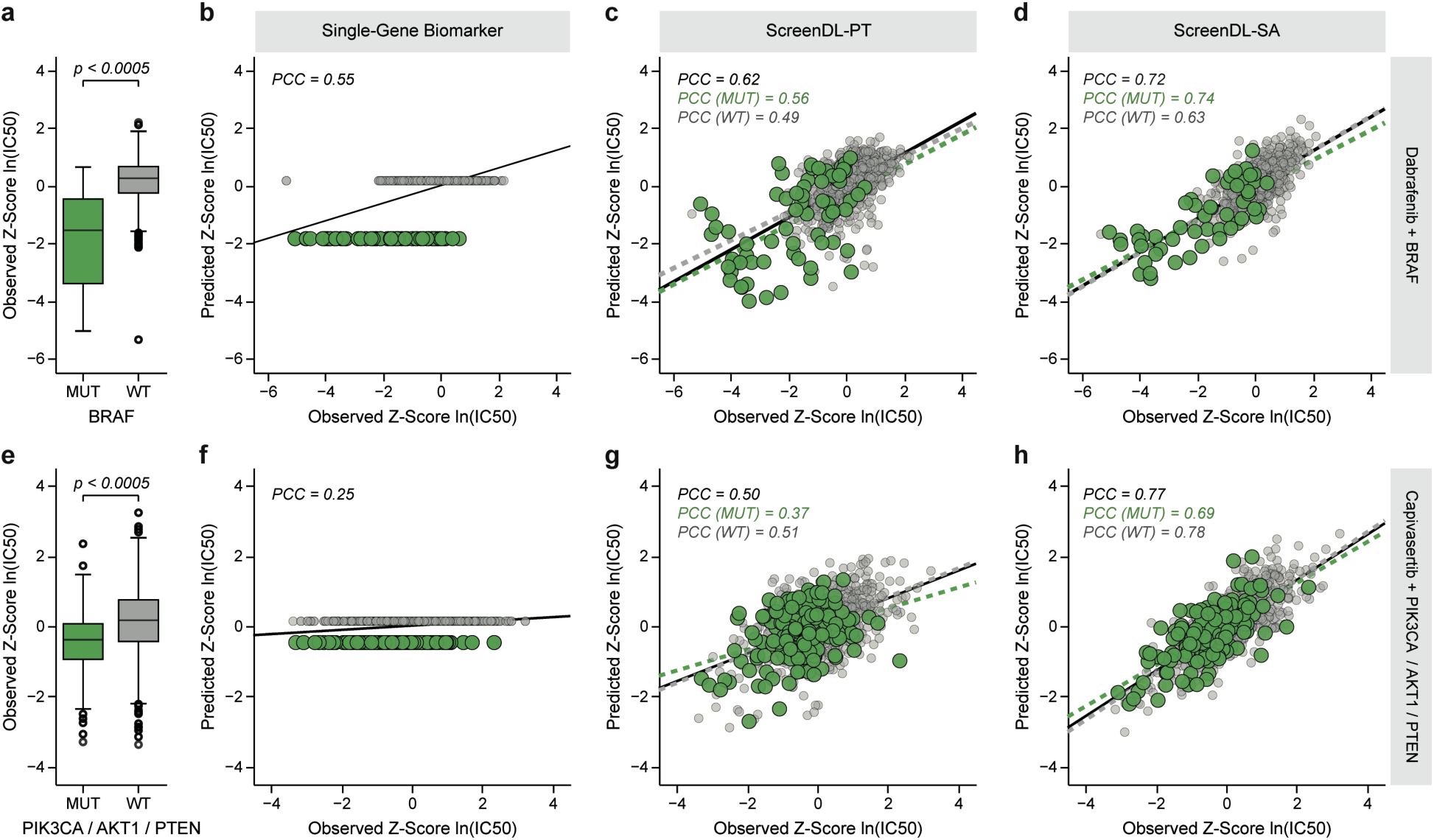
ScreenDL outperforms single- and multi-gene biomarker-only models in cell lines. **a-d.** Sensitivity to the BRAF inhibitor dabrafenib in cell lines with and without BRAF mutations compared by a two-sided Mann-Whitney U test (**a**). Performance of a biomarker only-model (**b**) compared with ScreenDL-FT (**c**) and ScreenDL-SA (**d**). **b-d.** Lines indicate linear regressions fit to the data. **c-d.** Performance of ScreenDL-FT and ScreenDL-SA is stratified by genomic subgroups **e-h.** Sensitivity to the AKT inhibitor capivasertib in PDXOs with and without mutations in PI3KCA, AKT1, and/or PTEN compared by a two-sided Mann-Whitney U test (**e**). Performance of a biomarker only-model (**f**) compared with ScreenDL-FT (**g**) and ScreenDL-SA (**h**). **f-h.** Lines indicate linear regressions fit to the data. **g-h.** Performance of ScreenDL-FT and ScreenDL-SA is stratified by genomic subgroups

Replicating our cell line analysis, we also observed enhanced capivasertib sensitivity in breast cancer PDXOs harboring *PIK3CA, AKT1*, and/or *PTEN* mutations (p = 0.02, Mann-Whitney U test; Fig. 5a). Again, ScreenDL-SA outperformed a biomarker-only model of capivasertib sensitivity, achieving a PCC of 0.50 compared to 0.46 (Fig. 5b-d). ScreenDL-SA also successfully identified the capivasertib-sensitive line, BCM-2665, representing a patient lacking the qualifying driver mutations who might benefit from capivasertib treatment. Similarly, ScreenDL-SA successfully identified BCM-5998 as capavisertib-resistant despite harboring a deleterious mutation in *PTEN*. We next focused on *BRCA1/2* mutations given their approval as a biomarker for response to PARP inhibition and their association with sensitivity to platinum-based chemotherapy.^49^ As expected, deleterious *BRCA1/2* mutations were associated with heightened sensitivity to the PARP1/2 inhibitor talazoparib (p = 0.03, Mann-Whitney U test; Fig. 5e) and the platinum-based chemotherapy carboplatin in PDXOs (p = 0.01, Mann-Whitney U test; Fig. 5i). ScreenDL-SA outperformed *BRCA1/2* biomarker-only predictive models for both talazoparib and carboplatin while gaining power to stratify sensitive and resistant PDXOs within genomic subgroups. Further, both ScreenDL-FT and ScreenDL-SA uncovered a subpopulation of PDXOs with poor talazoparib response despite harboring deleterious *BRCA1/2* mutations (Fig. 5f-h). Similarly, ScreenDL-SA correctly identified several PDXO lines displaying above-average carboplatin sensitivity despite lacking detectable *BRCA1/2* alterations (Fig. 5l).

**Fig. 5:**
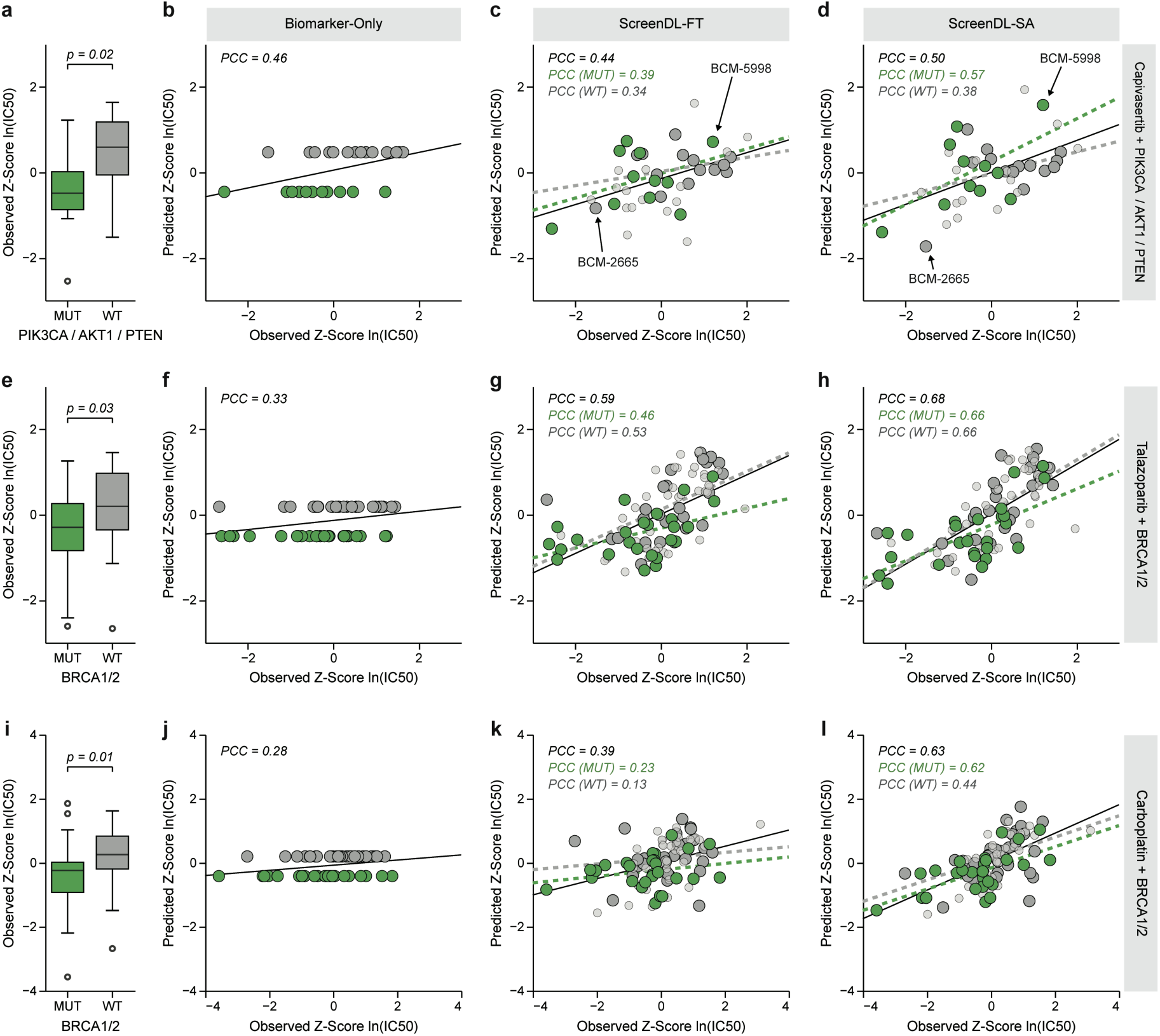
ScreenDL outperforms single- and multi-gene biomarker-only models in advanced/metastatic breast cancer PDXOs. **a-d.** Sensitivity to the AKT inhibitor capivasertib in breast cancer PDXOs with and without mutations in PI3KCA, AKT1, and/or PTEN compared by a two-sided Mann-Whitney U test (**a**). Performance of a biomarker only-model (**b**) compared with ScreenDL-FT (**c**) and ScreenDL-SA (**d**). **c-d.** Performance of ScreenDL-FT and ScreenDL-SA is stratified by genomic subgroups. **e-h.** Sensitivity to the PARP inhibitor talazoparib in PDXOs with and without mutations in BRCA1/2 compared by a two-sided Mann-Whitney U test (**e**). Performance of a biomarker only-model (**f**) compared with ScreenDL-FT (**g**) and ScreenDL-SA (**h**). **g-h.** Performance of ScreenDL-FT and ScreenDL-SA is stratified by genomic subgroups. **i-l.** Carboplatin sensitivity in PDXOs with and without mutations in BRCA1/2 compared by a two-sided Mann-Whitney U test (**e**). Performance of a biomarker only-model (**j**) compared with ScreenDL-FT (**k**) and ScreenDL-SA (**l**). **k-l.** Performance of ScreenDL-FT and ScreenDL-SA is stratified by genomic subgroups. **b-d,f-h,j-l.** Lines indicate linear regressions fit to the data. **c-d,g-h,k-l.** PDXOs without whole exome sequencing (WES) data are indicated in light gray.

### ScreenDL consistently selects efficacious therapies for breast cancer PDX models

The relationship between PDX and PDXO mirrors that of patient and PDO, affording an opportunity for end-to-end validation of our treatment selection strategy in a preclinical system that parallels clinical application (Fig. 6a). To this end, we applied our combined computational/experimental approach retrospectively to 20 breast cancer PDX models for which sufficient *in vivo* response data was available. For each PDX, bulk RNA-seq of the PDX’s tumor (or the PDXO when PDX sequencing was not available) and partial drug screening (n=12) from matched PDXOs were integrated into ScreenDL through ScreenAhead, and the drug with the lowest predicted *Z_D_* response from a panel of candidate therapies was selected as the optimal precision treatment (Methods; Fig. 6a). In total, this panel contained between two and nine drugs for each PDX line. Following the recommendations of Meric-Bernstam and colleagues^50^, we quantified performance in terms of: (1) clinical benefit rate (CBR), defined as the fraction of PDX lines displaying stable disease (SD) or better by mRECIST criteria^20^; and (2) objective response rate (ORR), defined as the fraction of PDX lines achieving at least a partial response (PR). ScreenDL-SA achieved an 85% CBR and a 55% ORR (Fig. 6d,g), with 8 of 20 PDX lines achieving a complete response (CR) (Fig. 6h). In comparison, drug selection using the raw PDXO screening data achieved a 75% CBR and a 35% ORR (Fig. 6b,e). Notably, 4 of 5 HCI-019 mice achieved an over 99% reduction in tumor volume after talazoparib therapy despite receiving an mRECIST designation of stable disease (SD) due to a weak initial response. When reclassifying the response of HCI-019 to talazoparib as a CR, ScreenDL-SA achieved an improved ORR of 60%. Further, while drug selection with ScreenDL-SA significantly improved ORR (p = 0.006, Fisher exact test; Fig. 6h,i), we did not observe significant improvement when selecting drugs using the raw PDXO screening data (p = 0.33, Fisher exact test). As a whole, these results are highly encouraging, particularly given evidence that a strong response in PDX models is more predictive of clinical benefit in patients.^50^

**Fig. 6:**
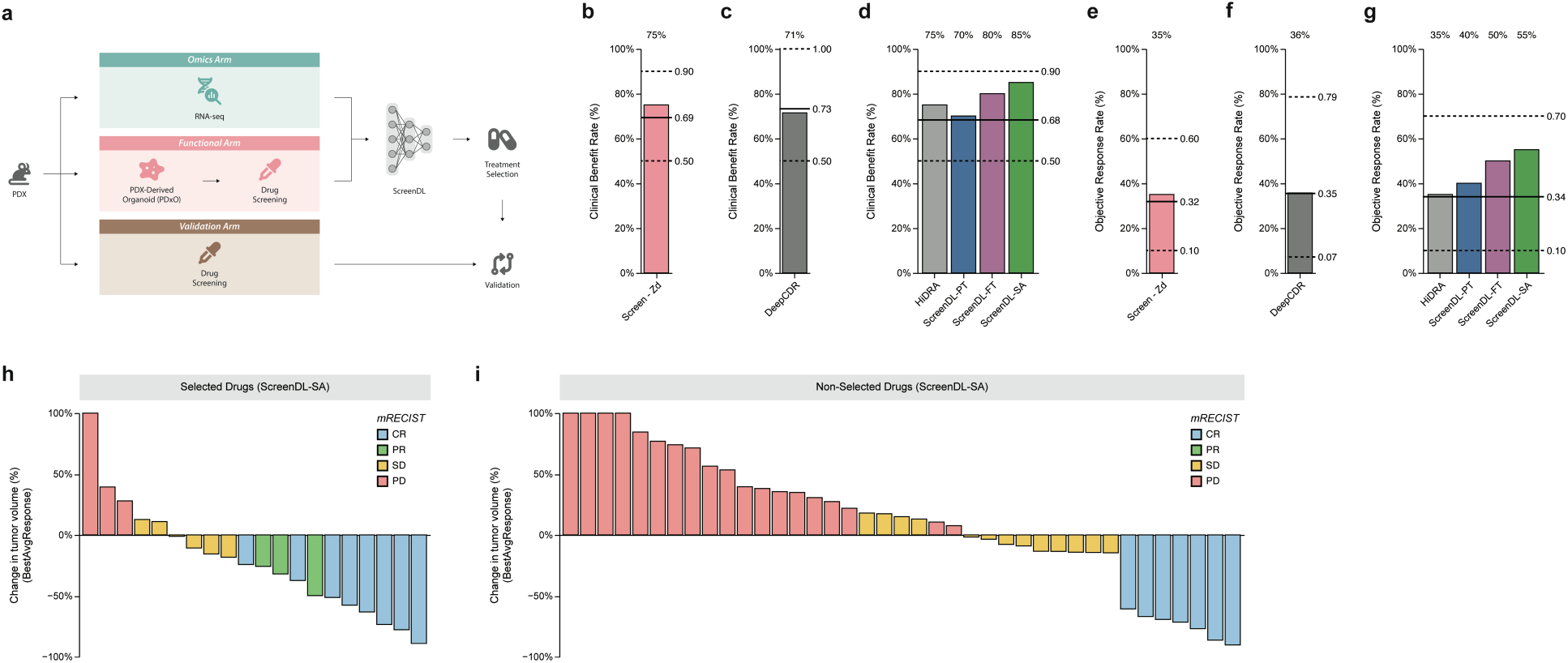
Retrospective validation of our end-to-end precision treatment selection strategy in matched PDX/PDXO models. **a.** Retrospective Validation Schema. A PDX’s transcriptomic profile and functional drug screening from matched PDXO lines are integrated into ScreenDL through tumor-specific fine-tuning with ScreenAhead. For each PDX, the drug with the lowest predicted *Z_D_* is selected as the optimal treatment and both clinical benefit rate (CBR) and objective response rate (ORR) are quantified using *in vivo* response in the originating PDX lines. Only the 20 PDX models for which *in vivo* testing for at least two therapies was carried out in unrelated studies were considered. **b.** CBR amongst the drugs selected with raw PDXO screening data for 20 PDX lines. Response values were *z*-score normalized independently for each drug across tumor samples and the drug with the lowest *z*-score response in the corresponding PDXO line was selected as the optimal precision treatment. Clinical benefit was defined as stable disease (SD) or better by mRECIST criteria. **c.** CBR amongst drugs selected by DeepCDR for the subset of 15 PDX lines with WES data. **d.** CBR for drugs selected by each ScreenDL variant for 20 PDX lines compared with those selected by HiDRA. **e.** ORR amongst the drugs selected with raw PDXO screening data for 20 PDX lines. Objective response was defined as partial or complete response (PR or CR) by mRECIST criteria. **f.** ORR amongst drugs selected by DeepCDR for the subset of 15 PDX lines with WES data. **g.** ORR amongst the drugs selected by each ScreenDL variant for 20 PDX lines compared with those selected by HiDRA. **b-g.** Solid lines correspond to the CBR (**b-d**) or ORR (**e-g**) achieved by random drug selection. Dashed lines correspond to the maximum and minimum achievable CBR (**b-d**) or ORR (**e-g**) based on the observed PDX screening data. PDXO screening was not performed for a subset of drugs evaluated in PDX lines, resulting in changes in the CBR/ORR achieved by random drug selection, as well as the minimum/maximum achievable CBR/ORR when using either DL models or raw PDXO screening data. **h.** Waterfall plot showing changes in tumor volume quantified as the mean BestAvgResponse across mice (see Methods) for PDX lines treated with optimal precision therapies selected by ScreenDL-SA. **i.** Waterfall plot showing changes in tumor volume quantified as the mean BestAvgResponse across mice for PDX lines treated with drugs that were not selected as optimal precision therapies by ScreenDL-SA. **h,i.** Color indicates mRECIST classifications for each PDX-drug pair. Positive changes in tumor volume are capped at 100%.

## Discussion

We have developed a unified computational/experimental approach to precision treatment selection, coupling a comprehensive patient-derived tumor model characterization and functional testing pipeline with ScreenDL, a DL-based CDRP framework engineered to exploit the highly valuable data generated by this experimental workflow. In particular, ScreenDL leverages data generated by this experimental arm, first for domain-specific fine-tuning with our expanded PDXO pharmaco-omic dataset and second, for personalization through tumor-specific fine-tuning with ScreenAhead. At baseline, our pretrained model displays enhanced capabilities of extrapolating to unseen tumor samples and outperforms existing DL methods on a series of precision oncology-relevant benchmarks. While current precision oncology practice is limited to targeted agents, ScreenDL provides accurate predictions for both chemotherapies and targeted agents with diverse biological mechanisms, facilitating selection from the full spectrum of anticancer therapies. This work establishes a unified framework for the integration of tumor omic and functional data from the same originating tumor sample through DL, powering ScreenDL for therapy selection and bringing AI-guided precision oncology into the realm of clinical application.

The superior performance of ScreenDL in PDMCs is driven by the wealth of domain-specific pharmaco-omic data generated by our experimental tumor characterization pipeline. This rich dataset links tumor omic features with drug response phenotypes in 100 breast cancer PDXO/PDO models derived primarily from high-risk, endocrine resistant, treatment-refractory, and/or metastatic tumors representing the greatest unmet clinical and research needs in breast cancer care. Characterization of this pharmaco-omic resource revealed strong concordance of functional drug sensitivities with known genomic anchors. Further, *in vivo* validation of selected responses revealed that PDXO screening was strongly predictive of *in vivo* response in the originating PDX model, highlighting the ability of our PDXO system to recapitulate many of the complexities of drug response in human tumors. Along with the computational methods described herein, we make this dataset available through cBioPortal as a public resource for use by the research community.

We have introduced ScreenAhead, a patient-specific fine-tuning strategy capable of exploiting paired omic and functional data from the same originating tumor sample to generate personalized response predictions. While a number of DL-based CDRP methods have been developed in recent years^14^, the ScreenAhead approach described here represents, to our knowledge, the first method enabling the direct incorporation of tumor omic and functional data from the patient-of-interest into a DL-based CDRP algorithm. We have shown that ScreenAhead effectively exploits functional patterns in partial screening data to circumvent the limitations of omics alone for drug response inference. In particular, ScreenAhead derives knowledge of global drug sensitivity (GDS) from partial screening data and exploits learned drug-drug functional similarities to improve predictions for unscreened drugs. As a result, ScreenDL-SA provides high-confidence predictions for chemotherapies and targeted agents with diverse biological mechanisms. Further, by requiring only a limited pre-screening panel, our ScreenAhead strategy has the potential to significantly reduce the overhead of patient-specific functional testing while still exploiting the ability of functional data to augment omic information and enhance treatment selection. Paired with our experimental protocol capable of returning tumor omic and functional drug screening results within a clinically relevant timeframe, the superior performance afforded by ScreenAhead represents a significant advance in DL-based CDRP.

Here, the superior predictive power of ScreenDL-SA warrants discussion of the broader clinical feasibility of patient-specific functional testing. Recent years have seen the development of organoid screening platforms and other short-term *ex vivo* culture systems maintaining strong biological fidelity and concordant drug responses with tumors from diverse cancer types.^23,29,51–54^ Collectively, these and other advancements have motivated numerous ongoing clinical trials seeking to exploit *ex vivo* functional testing for personalized therapy selection, including evaluation of the hybrid computational/experimental treatment selection protocol described here in our Functional Precision Oncology for Metastatic Breast Cancer study (FORESEE; ClinicalTrials.gov: NCT04450706).^29^ Grounded by these ongoing efforts to incorporate functional testing into clinical decision-making, ScreenAhead represents a powerful tool for the integration of patient-specific omic and functional data to enhance precision treatment selection.

We have demonstrated the clinical potential of our end-to-end treatment selection framework with a preclinical pilot in breast cancer PDX models. Here, the dichotomy between PDX and PDXO parallels clinical application in patient tumors with functional testing in matched PDOs, providing a surrogate assessment of clinical utility. Therapies selected by ScreenDL-SA achieved an 85% CBR and a 55% ORR, outperforming those selected by existing DL models or raw PDXO screening. Here, ScreenDL’s superior treatment selection capabilities relative to raw PDXO screening alone demonstrate the benefit of incorporating pharmaco-omic data from cell lines and PDXOs in a pretraining/fine-tuning schema. We note that a subset of PDX-drug pairs in this validation was selected for *in vivo* testing, at least in part, due to treatment efficacy in the corresponding PDXO lines. As such, this retrospective validation may overstate the CBR and ORR in unselected populations. Despite this limitation, we anticipate improved treatment selection in future prospective validations, as the retrospective study described here limits candidate treatments to those already tested in the corresponding PDX for unrelated studies. Thus, the selected treatments may represent suboptimal therapies when considering a larger panel of candidate agents. Future prospective validations wherein the best predicted treatment is chosen from a panel of high-confidence drugs for which ScreenDL provides accurate predictions in PDXOs will likely yield further improvements in ScreenDL’s treatment selection capabilities.

Despite promising results in this preclinical validation, our treatment selection framework is not without limitations. Beyond the need for patient-specific functional testing for ScreenAhead, a large resource of cancer type-specific PDMC pharmaco-omic data for domain-specific fine-tuning is necessary for optimal performance (Extended Data Fig. 7). Nonetheless, our breast cancer application demonstrates the power of such tumor type-specific PDMC resources for fine-tuning ScreenDL. While costly, we anticipate that the future generation of comparable PDMC resources will power ScreenDL for application in diverse cancer types. In addition, as with most PDMCs, our organoid platform lacks key elements of the tumor microenvironment, namely immune and stromal cells, preventing application of our approach for immunotherapy selection^21^.

This work establishes a unified computational/experimental approach to precision treatment selection in high-risk/metastatic breast tumors, pairing advances in *ex vivo* functional testing with a computational pipeline designed to integrate patient-specific omic and functional drug screening data to provide accurate response predictions tailored to individual tumors. This approach consistently selects efficacious therapies in preclinical validations which closely approximate clinical application, justifying further prospective preclinical studies and laying the groundwork for evaluation in future clinical trials.

## Methods

### Pharmaco-omic data harmonization and feature extraction

To establish an extensive pharmaco-omic dataset for pretraining, drug response data was retrieved from the Genomics of Drug Sensitivity in Cancer (GDSC) database^34^ and integrated with omics data from Cell Model Passports^35^. To facilitate comparison with existing DL models requiring multi-omic inputs, only cell lines with complete multi-omic profiles including mutations, transcriptomics, and copy number profiles were considered, ultimately yielding a harmonized pretraining dataset consisting of 278,033 tumor-drug pairs spanning 799 pan-cancer cell lines and 409 anticancer therapies. Drug response was quantified using GDSC-provided IC50 values, the most common dose-response metric used in DL-based CDRP models^14^. To standardize dose-response metrics across GDSC releases (i.e., GDSC1 and GDSC2), raw IC50 values were natural log (ln) transformed and *z*-score normalized independently for each drug. For drugs included in both GDSC1 and GDSC2, only drug responses from GDSC2 were retained to remove technical artifacts arising from the particular dose-response assay used in each release. For each compound in the resulting dataset, canonical SMILES strings were queried from PubCHEM using GDSC-provided PubCHEM^55^ compound identifiers (CIDs). CIDs were manually curated for drugs lacking GDSC-provided CIDs. Morgan fingerprints (512 bits, radius = 3) were generated from the canonical SMILES notion of each drug using RDKit (https://www.rdkit.org). To generate computational representations of each tumor, raw TPM gene expression values for 4,364 genes in the Molecular Signatures Database hallmark gene set collection^32^ were log_2_ transformed (with a pseudocount of 1). The resulting values were *z*-score normalized across tumor samples, providing vectors of transcriptomic features representing well-defined biological states and cellular processes for response prediction.

Omic data generation for PDX/PDXO samples is detailed below. In what follows, we focus on the harmonization of PDX/PDXO data with that of our cell line pharmaco-omic dataset. Raw TPM values from PDX/PDXO samples were log_2_ transformed and technical biases between cell line and PDX/PDXO expression were corrected using pyComBat^56^ prior to *z*-score normalization. IC50 response values for each PDXO-drug pair were generated using the GRmetrics R package^57^ and natural log (ln) transformed. Further detail regarding the calculation of dose-response curve metrics is provided in Supplementary Text 4. To maximize the PDXO response data available for fine-tuning, undefined ln(IC50) values were imputed using alternative dose-response metrics computed by the GRmetrics package. The resulting ln(IC50) values were *z*-score normalized independently for each drug across PDXO samples. We note that *z*-score normalization was performed separately for cell lines and PDXO samples to account for technical variations in dose-response measurements across domains. Ultimately, this yielded a PDXO dataset consisting of 4,278 drug responses spanning 98 PDXO lines and 107 anti-cancer agents.

### Model architecture & pretraining

ScreenDL consists of three full-connected subnetworks: (1) a **tumor omic subnetwork** which maps an input vector encoding 4,366 gene expression values to a reduced dimensionality representation (embedding) of a tumor’s transcriptomic profile; (2) a **drug subnetwork** which maps a drug’s 512 bit Morgan fingerprint to a reduced dimensionality embedding of chemical structure; and (3) a **shared response prediction subnetwork** which takes the concatenated tumor and drug embeddings as input and produces a predicted *Z_D_* value, corresponding to the z-score normalized, natural log transformed half-maximal inhibitory concentration or *z*-score ln(IC50) of a given tumor-drug pair. This multi-input framework in which independent tumor and drug subnetworks are dedicated to learning rich embeddings of transcriptomic and chemical features which are later integrated by a shared response prediction subnetwork represents the field-standard for DL-based CDRP models and was shown to outperform alternative configurations in a recent analysis.^14,58^ Each of ScreenDL’s subnetworks consisted of a series of fully connected layers configured as follows: (1) ScreenDL’s tumor-omic subnetwork was configured with four fully connected layers with 512, 256, 128, and 64 neurons; (2) ScreenDL’s drug subnetwork was configured with three fully connected layers with 256, 128, and 64 neurons; and (3) ScreenDL’s shared response prediction subnetwork was configured with two hidden layers with 128 and 64 neurons. Each fully connected layer was followed by Leaky ReLU activation with two exceptions: (1) the final output layer which was followed by a linear activation; and (2) the output layers of ScreenDL’s tumor omic and drug subnetworks which were followed by hyperbolic tangent activation. To help mitigate overfitting and enhance generalizability to never-before-seen tumor samples, a gaussian noise layer was added before the first fully connected layer of ScreenDL’s omics subnetwork. ScreenDL was implemented in Python using the Tensorflow/Keras framework and trained on GPU nodes provided by the Utah Center for High Performance Computing equipped with either NVIDIA GTX 1080 Ti GPUs with 3584 CUDA cores and 11 GB GDDR5X memory or NVIDIA A40 GPUs with 10,752 CUDA cores and 48 GB GDDR6 memory.

### Evaluation of ScreenDL in cancer cell lines

Evaluation in cell lines followed a 10-fold tumor-blind cross-validation schema with performance evaluated on never-before-seen cell lines, mirroring clinical application in never-before-seen patients. Specifically, each fold consisted of three disjoint sets of cell lines (and their corresponding drug response observations) used for training (*T*), validation (*V*), and testing/evaluation (*E*), respectively. Sample proportions were held constant across folds with each fold’s *T*, *V*, and *E* sets consisting of all drug response observations for 80%, 10%, and 10% of cell lines, respectively. Cell lines derived from different cancer types were evenly distributed across folds to ensure adequate representation of individual disease types in each training set. For each fold, the validation set *V* was used for early termination to prevent overfitting. ScreenDL-PT’s parameters were optimized to minimize the mean squared error between observed and predicted *Z_D_* response values using the Adam optimizer with a mini-batch size of 256, a learning rate of 0.0001, and weight decay set to 0.0001.

To provide a comprehensive assessment of precision oncology-relevant performance, we adopted a set of three performance metrics providing distinct yet complementary views of predictive power. As a primary assessment, we quantified prediction accuracy independently for each drug in terms of the Pearson correlation coefficient (PCC) between observed and predicted *Z_D_* values. In addition, we binarized observed *Z_D_* values and quantified area under the receiver operating characteristic curve (auROC) for each drug, providing a measure of ScreenDL’s power to stratify sensitive and resistant tumor samples. Finally, we applied ScreenDL to select the precision treatment that minimized the predicted *Z_D_* for each tumor sample and quantified response rate (RR) as the fraction of tumor samples for which observed response fell below the 30th percentile of *Z_D_* values for the selected therapy. While both PCC and auROC quantify drug-level performance, RR provides a binary measure of success for individual tumor samples and thus provides a direct assessment of treatment selection capabilities. Stratified analysis of drug level performance was performed using manually curated annotations of drug type (i.e., chemotherapy vs. targeted agent) for a subset of 213 therapies (34 chemotherapies and 179 targeted agents). For pathway-level analysis, we used GDSC-provided target pathway annotations augmented with manually curated annotations for drugs not included in the GDSC database.

ScreenAhead tumor-specific fine-tuning was performed independently for each cell line. Specifically, for each cell line, a subset of 20 drugs was selected for pre-screening using principal feature analysis (PFA). Here, we frame drug selection for ScreenAhead as a feature selection task on an *MxN* drug response matrix where each row corresponds to one of *M* cell lines in the training dataset and each column (feature) corresponds to one of *N* candidate drugs. We applied PFA to select the subset of these *N* drugs that described most of the variation in observed drug responses across cell lines in the training dataset. For each cell line, we fine-tuned ScreenDL-PT for 20 epochs using the cell line’s transcriptomic profile and its observed responses to the resulting set of 20 pre-screening drugs. During fine-tuning, ScreenDL-PT’s tumor omic and drug subnetworks were frozen and parameters in the shared response prediction subnetwork were updated using the Adam optimizer with a learning rate of 0.0001. The resulting ScreenDL-SA model was applied to predict response for the remaining unscreened drugs in a given cell line, and performance was assessed following the procedure outlined for ScreenDL-PT.

### Comparison of ScreenAhead drug selection algorithms

To ensure optimal performance with ScreenAhead, we evaluated three informed drug selection algorithms designed to maximize coverage of the drug response space and ensure systematic exploration of potential therapeutic responses. Specifically, we compared the performance of ScreenDL-SA in cell lines when selecting pre-screening drugs under two scenarios: random drug selection, and informed selection using either PFA, agglomerative clustering, or metadata annotations of drug biological mechanisms. Each drug selection algorithm was applied to select pre-screening sets of 5, 10, 15, 20, and 25 drugs for each cell line, and ScreenAhead tumor-specific fine-tuning was performed with each set as described above. For a given number of pre-screening drugs, pairwise statistical comparisons of each algorithm’s drug-level performance were performed with two-sided Wilcoxon signed-rank tests.

### Evaluation of ScreenDL in breast cancer PDXOs

We assessed the performance of ScreenDL in breast cancer PDXOs by training 10 ScreenDL-PT models on different subsets of cell lines. Specifically, cell lines were split into 10 disjoint sets, and 10 ScreenDL-PT models were trained on cell lines from 9 of these 10 sets (i.e., pharmaco-omic data from 90% of all cell lines). During pretraining, the remaining 10% of cell lines were used for early stopping. This produced an ensemble of ten ScreenDL-PT models which were each applied to predict treatment response in our PDXO cohort. For a given PDXO-drug pair, the trimmed mean of predicted *Z_D_* responses across these ten ensemble members was taken as the final prediction and performance was assessed for each of precision oncology-relevant metrics detailed above. To ensure fair comparisons with DeepCDR and HiDRA, the same ensembling procedure was used for all models. We note that DualGCN was not carried forward for evaluation in PDXOs due to its weak performance in cell lines and its requirement of copy number features.

To fine-tune ScreenDL for response prediction in breast cancer PDXOs, parameters in ScreenDL-PT’s drug subnetwork and those in the first hidden layer of ScreenDL-PT’s tumor omic subnetwork were frozen. Using our PDXO pharmaco-omic dataset, we fine-tuned ScreenDL-PT for 30 epochs using the Adam optimizer with a mini-batch size of 256, an initial learning rate of 0.0001, and weight decay set to 0.01. The learning rate was held constant during the first 2 epochs and decayed exponentially for each remaining epoch. Performance was assessed under a leave-one-out cross-validation schema wherein ScreenDL-PT was fine-tuned with pharmaco-omic data from all PDXOs except one and predictions were generated for the remaining left-out PDXO line. To prevent data leakage, PDXO lines derived from the same originating tumor sample as a given left-out PDXO were excluded from fine-tuning. Each of ScreenDL-PT’s ensemble members was fine-tuned independently, producing ten ScreenDL-FT models. Final predictions for each PDXO-drug pair were generated by taking the trimmed mean of predicted *Z_D_* values across ensemble members and performance was reported for each of the precision oncology-metrics detailed above.

We benchmarked ScreenAhead in breast cancer PDXOs following the same general procedure described for cell lines. In brief, after domain-specific fine-tuning, we performed ScreenAhead tumor-specific fine-tuning using a subset of 12 pre-screened drugs for each PDXO. Here, we opt to use 12 drugs for ScreenAhead in PDXOs rather than a 20-drug panel for two reasons: (1) 12 represents a feasible number of drugs to pre-screen in PDOs within a clinically relevant timeframe; and (2) ScreenAhead with just 10 drugs in cell lines provided only slightly weaker performance compared to ScreenAhead using a larger 20 drug panel (Extended Data Fig. 4a). ScreenAhead tumor-specific fine-tuning was performed independently for each ensemble member and final predictions were generated as described for ScreenDL-FT.

### Comparisons with existing DL models

Relative to other omic modalities, gene expression is known to provide superior predictive power in CDRP models^59,60^. Thus, we compared ScreenDL’s performance to that of three existing DL models that also incorporate transcriptomic features – DeepCDR, DualGCN, and HiDRA. For each model, the generation of tumor and drug features was performed as described in the original publications. Each model was fit using published hyperparameters and evaluated under the same 10-fold tumor-blind cross-validation schema described for ScreenDL. We note that DeepCDR was only evaluated on a subset (n = 61) PDXO lines for which whole exome sequencing (WES) was available for the extraction of mutation-based features. Statistical comparisons of each model’s drug-level performance were performed with two-sided Wilcoxon signed-rank tests.

### Comparisons with biomarker-only models

To compare ScreenDL’s performance with that of approved single- and multi-gene biomarkers, we generated biomarker-only predictive models in both cell lines and PDXOs. Cell lines were designated as either mutant (MUT) or wild-type (WT) based on driver mutation status extracted from annotated VCF files provided by Cell Model Passports. In PDXOs, MUT/WT status was assigned from WES data using our published variant filtration criteria^21^. The statistical significance of each drug-biomarker combination was assessed with a Mann-Whitney U test comparing the observed *Z_D_* responses between MUT and WT tumors. Biomarker-only models were defined as binary conditional functions returning the mean observed *Z_D_* in MUT lines for tumors harboring the corresponding biomarker and the mean observed *Z_D_* in WT lines otherwise. ScreenDL-PT and ScreenDL-SA were trained following the procedure outlined above for either cell lines or PDXOs with the exception that, for a given drug-biomarker combination, the drug of interest was excluded from ScreenAhead tumor-specific fine-tuning. For both ScreenDL-PT and ScreenDL-SA, performance was assessed by computing the PCC between observed and predicted *Z_D_* across all tumors and within WT and MUT subgroups.

### Validation in matched PDX/PDXO models

To validate ScreenDL’s treatment selection capabilities, we selected a subset of PDX/PDXO pairs for which sufficient *in vivo* PDX response data had been collected for unrelated studies. PDX/PDXO pairs were retained for analysis if at least two drugs were tested in the corresponding PDX line, yielding a set of 20 PDX/PDXO pairs for validation. Only drugs that showed efficacy in at least one PDX line were considered. Ultimately, each PDX had been screened with between two and nine candidate agents. ScreenDL was trained as described above, with the exception that performance on the remaining n = 78 PDXO lines was used as an early termination condition during cell line pretraining. For each model, the drug with the lowest predicted *Z_d_* for a given PDX line was selected as the optimal precision treatment. Observed changes in PDX tumor volume were classified using the modified RECIST criteria (mRECIST) detailed by Gao and colleagues^20^ and performance was assessed in terms of: (1) clinical benefit rate (CBR), defined as the fraction of PDX lines with at least stable disease (SD) after treatment with the selected agent; and (2) objective response rate (ORR), defined as the fraction of PDX lines displaying either a partial or a complete response after treatment with the selected agent. We note that DeepCDR was assessed on the subset of 15 PDX lines for which WES data was available. To compare ScreenDL’s treatment selection capabilities with those of raw PDXO screening, we also used the raw *Z_d_* values from our PDXO drug screening experiments to select precision therapies for each PDX line. Baselines for random treatment selection were obtained by performing 1,000 iterations of random drug selection for each PDX line and taking the mean CBR/ORR across iterations. Fisher exact tests were used to compare CBR and ORR for selected vs unselected PDX-drug pairs.

### PDXO culture and drug treatment

PDXOs were established and cultured as previously described.^21,39^ Each PDXO line was validated to be composed of human tumor cells matching their source PDX and patient samples by STR analysis. Briefly, PDXOs were maintained in 6-well plates (Genesee Scientific, El Cajon, CA, USA) and cultured in 200-μl Matrigel (Corning, Corning, NY, USA) domes within Advanced DMEM/F12 (Thermo Fisher, Waltham, MA, USA) supplemented with 5% FBS, 10 mM HEPES (Thermo Fisher), 1× Glutamax (Thermo Fisher), 1 μg/ml hydrocortisone (Sigma-Aldrich, Burlington, MA, USA), 50 μg/ml gentamicin (Genesee Scientific), 10 ng/ml hEGF (Sigma-Aldrich), and 10 μM Y-27632 (Selleck Chemicals, Houston, TX, USA).

Mature organoids were collected using 80% dispase (Fisher Scientific, Waltham, MA, USA), 20% FBS, and 10 μM Y-27632 treatment (40U per well) at 37°C for 20 minutes. Approximately 40-50 mature organoids were seeded per well in 384-well tissue culture plates (PerkinElmer, Waltham, MA, USA), each with a solidified 10-μl Matrigel base layer and 30 μl of PDXO culture medium containing 5% Matrigel. A separate 384-well plate was seeded in 2 columns for day 0 normalization. Plates were incubated at 37°C and 5% CO2 overnight to allow organoids to settle onto the Matrigel base. After 24 hours, 30 μl of medium and 15 μl of CellTiterGlo 3D (Promega, Madison, WI, USA) were added to the day 0 plates. The plates were then incubated on a plate shaker (Benchmark Scientific, Sayreville, NJ, USA) at 500 rpm for 20 minutes and read using the EnVision XCite plate reader (PerkinElmer). A separate 1-ml deep 96-well drug plate (USA Scientific, Ocala, FL, USA) was prepared starting at 50uM with an eight-point 5x serial dilution, and 30 μl of each condition, in technical quadruplicate, was transferred to the seeded 384-well plates using the ViaFlo electronic 96-channel handheld pipette (Integra Biosciences, NH, USA). The dosed PDXO plates were covered with Breathe-Easy seals (USA Scientific) and incubated for 144 hours at 37°C and 5% CO2. After incubation, the seals were removed, and 15 μl of CellTiterGlo 3D was added to each well. The plates were incubated on a plate shaker at 500 rpm for 20 minutes and assayed using the EnVision XCite plate reader. Raw luminescence values from each condition were divided by those from the day 0 plate to calculate the fold change.

25 unique PDXO lines were screened in each of four phases (screens A-D; 100 lines total); 40-50 drugs were screened in each phase. In all four phases, three biological replicates were performed for each PDXO-drug pair to ensure data quality. Between phases, drug responses over the technical and biological replicates were analyzed and drugs were either carried forward to the next phase or dropped from future screening. Reasons for drug dropout included lack of availability, lack of any sign of efficacy across 25 models, or overt toxicity across 25 models. New drugs were also added at each phase as they became available.

### PDX transplantation and drug treatment

PDX transplantation procedures were performed as previously described^61,62^. Briefly, thawed PDX tumor fragments were implanted into the cleared mammary fat pad of 6-8-week-old female NRG mice (Jackson Laboratory stock 7799). Tumor sizes were measured twice weekly. Established tumors (∼100 mm³) were treated with eribulin (0.25mg/kg, IV, once per week) (Selleckchem, S8912), tolinapant (20mg/kg, oral gavage, seven days on/seven days off) (Selleckchem, S8681), birinapant (20mg/kg, IP, 3 doses/week) (NIH CTEP/CTP #756502), talazoparib (0.33mg/kg, oral gavage daily, 5 doses/week) (Selleckchem, S7048), doxorubicin (0.5mg/kg, IP, two cycles of day 1, day 8, day 15) (in house pharmacy), paclitaxel (10mg/kg, IP, day 28, once) (Selleckchem, S1150), SN-38/irinotecan (12.5mg/kg, oral gavage, 5 doses/week, 2 week on following one week off) (Selleckchem, S4908), alpelisib (30mg/kg, oral gavage, daily) (Selleckchem, S2814), RO4929097 (10mg/kg, oral gavage, 5 days/week) (Selleckchem, S1575), gemcitabine (50 mg/kg, IP, twice a week) (Selleckchem, S1714), olaparib (50mg/kg, oral gavage, daily) (Selleckchem, S1060), ponatinib (30mg/kg, oral gavage, daily) (Selleckchem, S1490), lapatinib (200mg/kg, oral gavage, daily) (Selleckchem, S2111), palbociclib (100mg/kg, oral gavage, 5 doses/week) (Selleckchem, S1579), AZD8186 (30 mg/kg, oral gavage for 21 days, using a 4-day-on, 3-day-off schedule) (Selleckchem, S7694), tamoxifen (20mg/kg, subQ, 5 doses/week) (Sigma Aldrich, T5648-1G), abemaciclib (75mg/kg, oral gavage, daily), (Selleckchem, S7158), vinorelbine (10mg/kg, IP, once per week) (Selleckchem, S4269), rapamycin (7.5mg/kg, IP, once per week) (Selleckchem, S1039), and everolimus (5mg/kg, oral gavage, 5 days per week), (Selleckchem, S1120). Final results were evaluated according to the modified RECIST (Response Evaluation Criteria In Solid Tumors)^63^ criteria (mRECIST) described by Gao and colleagues^20^.

### Omics for PDX and PDXO models

PDXO RNA and DNA was extracted as described previously^39^, and RNA and DNA from patient tumor samples were obtained from the Biorepository and Molecular Pathology/Molecular Diagnostics core. To extract RNA and DNA from PDX, 10-15 mg of flash frozen PDX tumor tissue was placed in a 2 ml safe-lock microtube (Biopur, #05-402-24C) and 450 μl of RLT Plus buffer with BME was added. After adding one 5 mm stainless steel bead (Qiagen #69989), the microtube was placed in a TissueLyser II (Qiagen) and lysed in three cycles or until lysate appeared clear (frequency 20/sec, duration 30 sec). Lysate was incubated for 10 min on ice and nucleic acids were isolated using the AllPrep micro kit (Qiagen # 80204) as described in Scherer et al.^39^ RNA and DNA concentrations were determined by Qubit broad range kits (Thermo Fisher #Q10211; #Q32850). All established patient-derived models were validated to match their original patient tumor using STR as described^39^.

Libraries for bulk RNA sequencing and whole exome sequencing (WES) were prepared and sequenced at the Huntsman Cancer Institute High-Throughput Genomics and Bioinformatics Core. For RNA-seq, lllumina TruSeq RNA Library Preparation kit v2 and the Illumina TruSeq Stranded Total RNA kit with Ribo-Zero Gold were used. For WES, Agilent SureSelectXT Human All Exon V6+ COSMIC, Agilent Human All Exon 50-Mb library or IDT xGEN Human Exome v2 with Nextera Flex library preparation protocols were used. WES and RNA-seq data was processed through the PDXNet pipelines^64,65^.

### Data processing for cBioPortal

For clinical data variables, meta data for samples was integrated with institutional patient data extracted from the electronic medical record using custom Python scripts and made both relative to the patient date of birth and formatted for cBioPortal (Dockerized version V6.0.16) per documentation (https://docs.cbioportal.org/file-formats). Variant Call Files (VCFs) from PDXNet pipelines were converted to Mutation Annotation Format (MAF) files using Ensembl Variant Effect Predictor VCF2MAF.pl. MAF files were compiled and formatted for cBioPortal using scripts available upon request. RNA-seq TPM results from PDXNet pipelines were merged by gene identifiers for analysis. Requisite *z*-score TPM values were calculated for all genes across samples using scripts provided by cBioPortal.org. All uploaded data passed cBioPortal integral validation scripts for expected formats and sample alignment.

### Data & Code Availability

All code used for model development and analysis is available at https://github.com/csederman/screendl. All data used for model training is downloadable from this repository or through cBioPortal (link to be provided upon acceptance for publication). ScreenDL is also available as a Singularity image through the IMPROVE (Innovative Methodologies and New Data for Predictive Oncology Model Evaluation) framework at https://github.com/JDACS4C-IMPROVE/Singularity.

## Supporting information

Supplementary Table S1. Summary of HCI breast cancer PDXO and PDX lines with clinical annotations

Supplementary Table S2. Summary of raw PDXO screening data

## Acknowledgements

We thank Lacey Dobrolecki for coordinating transfer of previously established BCM PDX to Huntsman Cancer Institute, and Tim Parnell for processing omics data through the established PDXNet pipelines. The computational resources used were partially funded by the NIH Shared Instrumentation Grant 1S10OD021644-01A1. This work was conducted with funding from the National Cancer Institute U54CA224076 and U01CA217617; the Breast Cancer Research Foundation Founders Fund; the Huntsman Cancer Institute Cancer Center Support Grant 3P30 CA042014-33; the Department of Defense Breast Cancer Research Program Breakthrough Award W81XWH1410417 and Era of Hope Scholar Award W81XWH1210077; the Huntsman Cancer Foundation; Baylor College of Medicine (BCM) CPRIT Core Facility Awards RP170691 and RP220646, as well as BCM P30 Cancer Center Support Grant NCI CA125123. Research reported in this publication utilized the University of Utah Genomics Core facility and the Huntsman Cancer Institute Biorepository and Molecular Pathology/Molecular Diagnostics, Preclinical Research Resource, High Throughput Genomics and Cancer Bioinformatics, and Research Informatics Shared Resources at Huntsman Cancer Institute at the University of Utah, supported by the National Cancer Institute of the National Institutes of Health under Award Number P30CA042014. The content is solely the responsibility of the authors and does not necessarily represent the official views of the NIH.

## Author Contributions

CS, TDS, and GTM conceived and designed the deep learning approach. CS, XH, YQ and GTM contributed to developing hypotheses and interpreting results. CS processed and harmonized the cell line and PDXO pharmaco-omic data. CS designed and architected the machine learning framework with feedback from TDS, GTM, and YQ. AW and BW oversaw PDX and PDXO establishment and AW, BW oversaw drug treatment experiments. LZ established PDXOs, ZC established PDXs and performed PDX transplantation and drug treatment. CHY and ECS performed PDXO culture and drug screens. ERW helped to maintain and expand the PDXO. SDS coordinated data generation and transfer. SDS, ERW, and CHY performed sample preparation for omic profiling. AA and JW processed omic data for PDMCs and uploaded them to cBioportal. AA and SDS processed metadata and developed a modified cBioportal instance. KTV and AW provided oversight to PDMC omics data sharing and cBioportal structure. MTL provided PDX models from BCM. CS, AW and GTM wrote the manuscript. All authors read and edited the manuscript.

## Extended Data Figures

**Extended Data Fig. 1:**
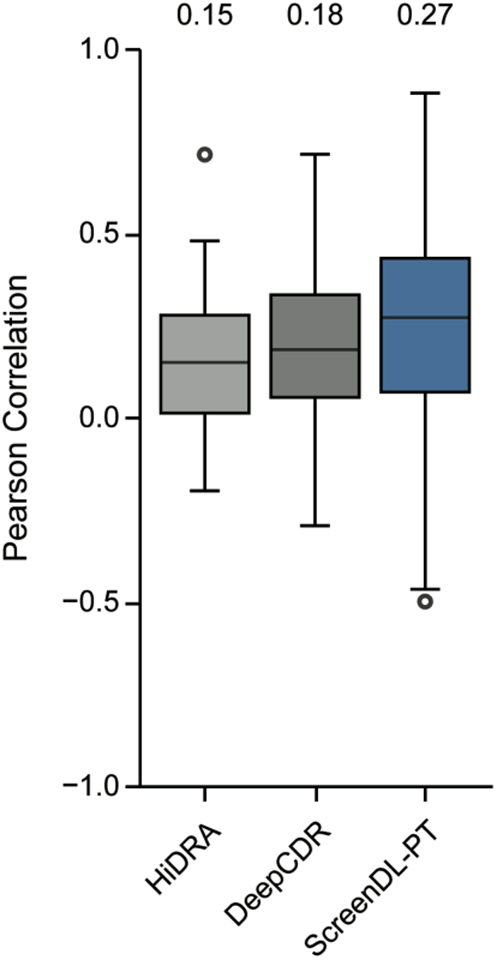
Performance in our previously reported collection of 16 high-risk/metastatic PDXO models. Performance comparison of ScreenDL-PT with two existing DL-based CDRP models, DeepCDR and HiDRA. Box plots represent the distribution of Pearson correlations between observed and predicted response per drug. Median values for each model are denoted in the upper margin.

**Extended Data Fig. 2:**
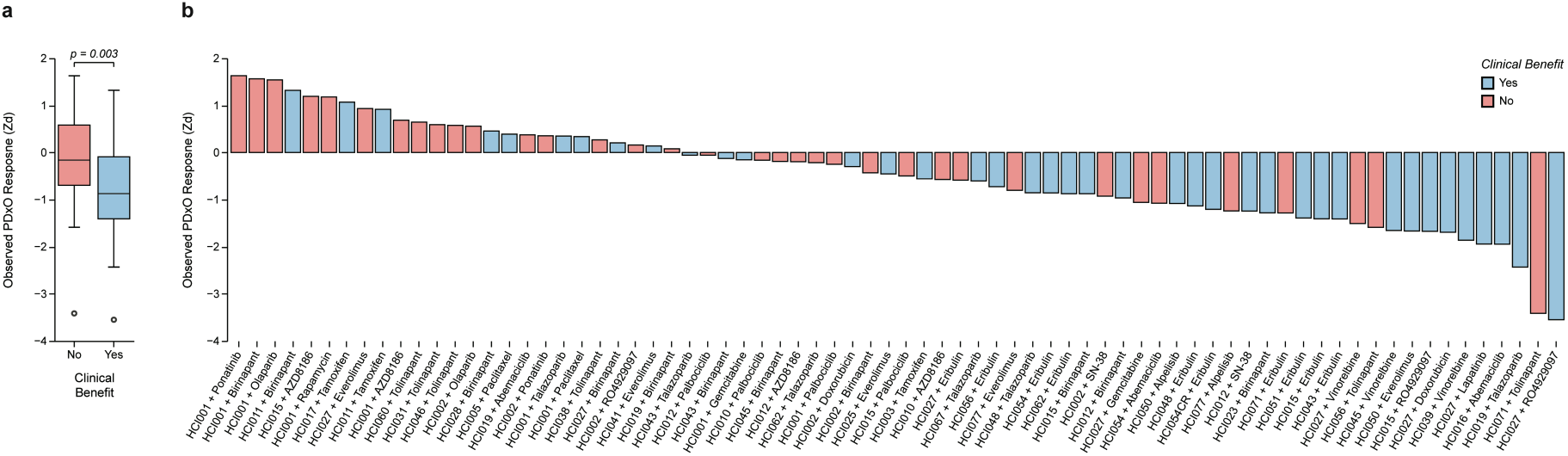
*In vitro* functional testing in PDXOs predicts *in vivo* PDX response. **a.** Boxplots of observed *Z_d_* response from raw PDXO screening stratified by whether or not the originating PDX line achieved clinical benefit after treatment with the corresponding therapy. Clinical benefit was defined as stable disease or better by mRECIST criteria. The p-value denotes the significance of a two-sided Mann-Whitney U test comparing observed *Z_d_* responses across groups. **b.** Bar graph representing observed *Z_d_* responses for the indicated drugs in PDXO lines. Color denotes whether or not the originating PDX showed clinical benefit after treatment with the corresponding therapy. An observed *Z_d_* below the 30th percentile of *Z_d_* values for a given therapy (corresponding to the top 30% most sensitive PDXOs for a given drug) was predictive of clinical benefit in the originating PDX line (p = 0.005, Fisher exact test).

**Extended Data Fig. 3:**
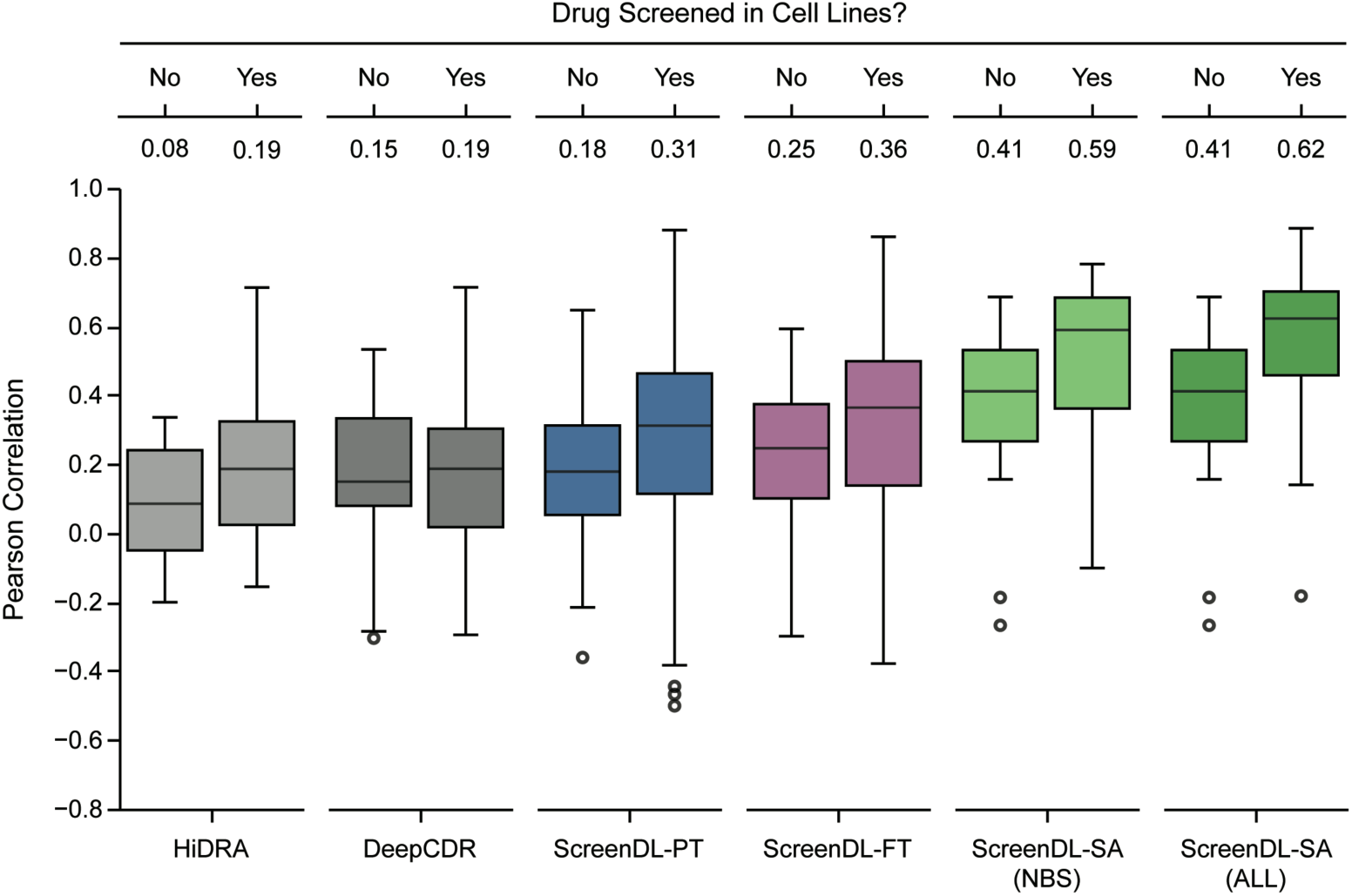
ScreenDL provides superior predictive power for drugs included in cell line pretraining. Drug-level performance of each ScreenDL variant and two existing DL models in breast cancer PDXOs stratified by whether or not drugs were included in cell line pretraining. Box plots represent the distribution of Pearson correlations between observed and predicted response per drug. Median values for each model are denoted in the upper margin. All models achieved superior performance for drugs included in cell line pretraining. We note that only drugs included in cell line pretraining were considered for ScreenAhead in PDXOs.

**Extended Data Fig. 4:**
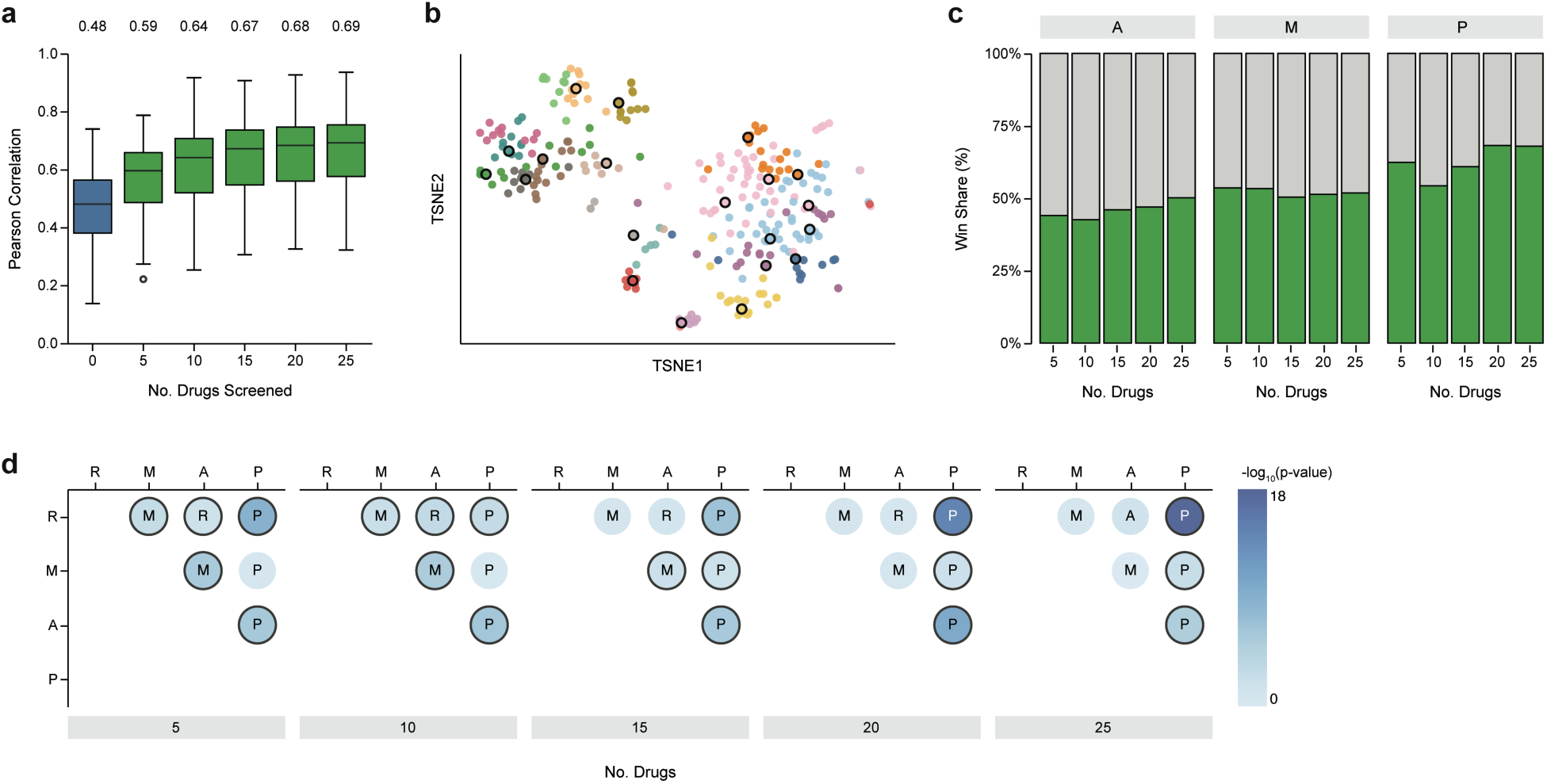
Systematic comparison of informed drug selection algorithms. **a.** Performance comparison of ScreenDL-PT (blue) and ScreenDL-SA (green) when adding an increasing number of randomly selected drugs to the ScreenAhead drug set. Box plots represent the distribution of Pearson correlations between observed and predicted response per drug. Median values are denoted in the upper margin. **b.** tSNE embeddings of drugs based on observed cell line responses. Colors indicate groups of drugs recovered by *k*-means clustering. Drugs selected for ScreenAhead using principal feature analysis are outlined in black. **c.** Percentage of drugs for which informed drug selection using either agglomerative clustering (A), metadata-based selection using drug target pathway annotations (M), or principal feature analysis (P) outperforms random selection. **d.** Pairwise comparison of informed drug selection algorithms. Per-drug performance was quantified as the Pearson correlation between observed and predicted response and drug-level performance was compared with Wilcoxon signed-rank tests. Circles are colored according to the negative log_10_ p-value and significant tests are outlined. The winner of each pairwise comparison is labeled within each circle.

**Extended Data Fig. 5:**
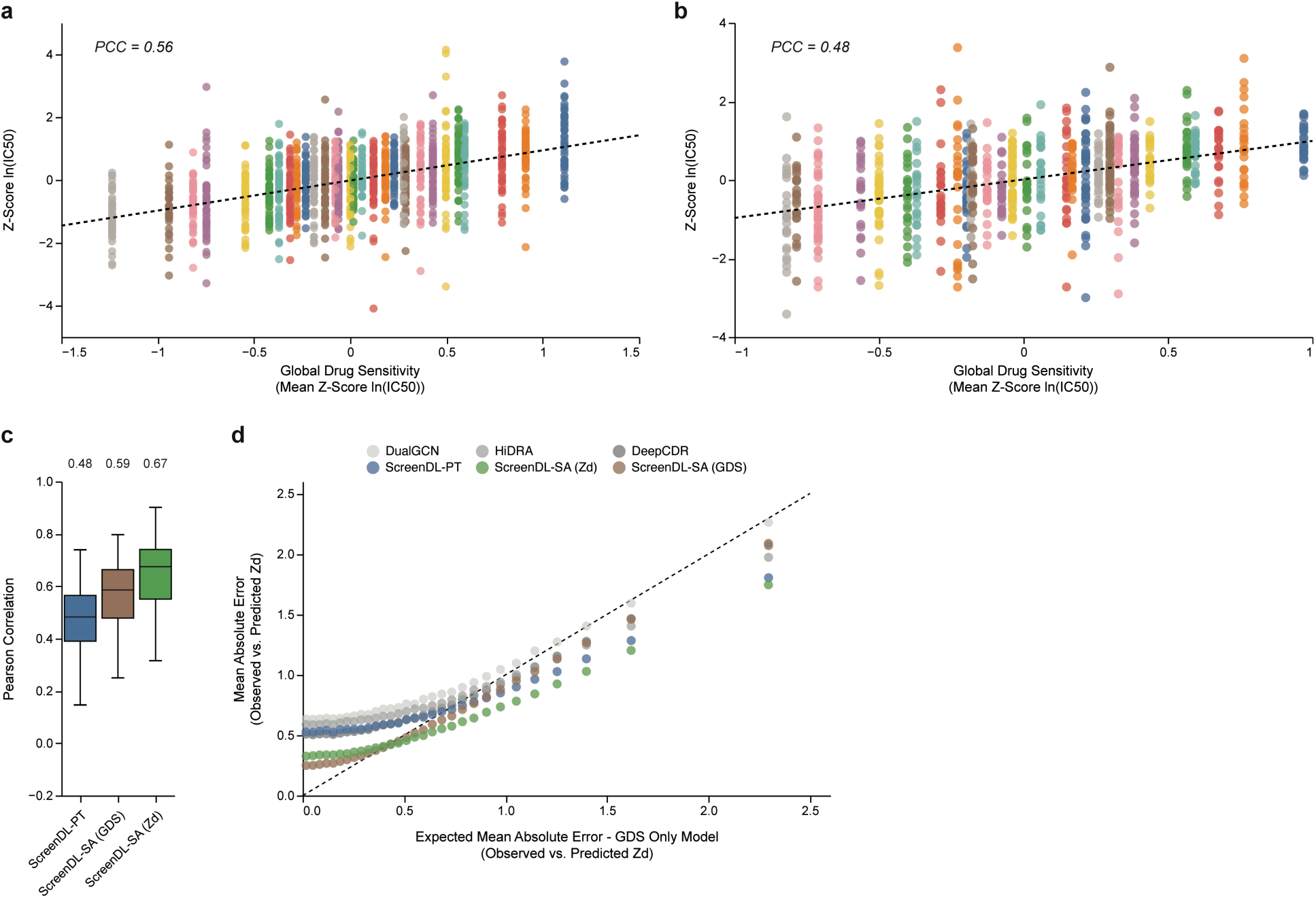
ScreenAhead leverages knowledge of global drug sensitivity encoded in partial pre-screening data. **a.** Correlation between global drug sensitivity (GDS) and response to individual therapies in cancer cell lines. For each cell line, GDS was defined as the mean *Z_d_* response across all screened drugs. Only a subset of 30 cell lines and 60 drugs are shown for readability. Colors indicate individual cell lines. The reported Pearson correlation represents the correlation between GDS and individual *Z_d_* values across the entire cell line dataset. **b.** Correlation between global drug sensitivity (GDS) and response to individual therapies in breast cancer PDXO models. For each PDXO, GDS was defined as the mean *Z_d_* response across all drugs. Only a subset of 30 PDXO lines and 60 drugs are shown for readability. Colors indicate individual PDXO lines. The reported Pearson correlation represents the correlation between GDS and individual *Z_d_* values across the entire PDXO dataset. **c.** ScreenDL-SA performance in cell lines when performing ScreenAhead tumor-specific fine-tuning using either the original *Z_d_* values (ScreenDL-SA (*Z_d_*)) or mean-filled *Z_d_* values for 20 drugs (ScreenDL-SA (GDS)). By replacing *Z_d_* values with a cell line’s mean *Z_d_* across the 20 ScreenAhead drugs, we effectively remove any drug-specific information and only provide knowledge of GDS during ScreenAhead tumor-specific fine-tuning. Box plots represent the distribution of Pearson correlations between observed and predicted response per drug. Median values for each ScreenDL variant are denoted in the upper margin. **d.** Mean absolute error (MAE) between observed and predicted *Z_d_* in cell lines for each ScreenDL variant and three existing DL models. The MAE of each model is binned by the expected MAE of a GDS-only model (i.e., a model that outputs a tumor’s GDS regardless of drug features; see Supplementary Text 2). ScreenAhead improves performance, even for cell line-drug pairs for which GDS is not highly predictive.

**Extended Data Fig. 6:**
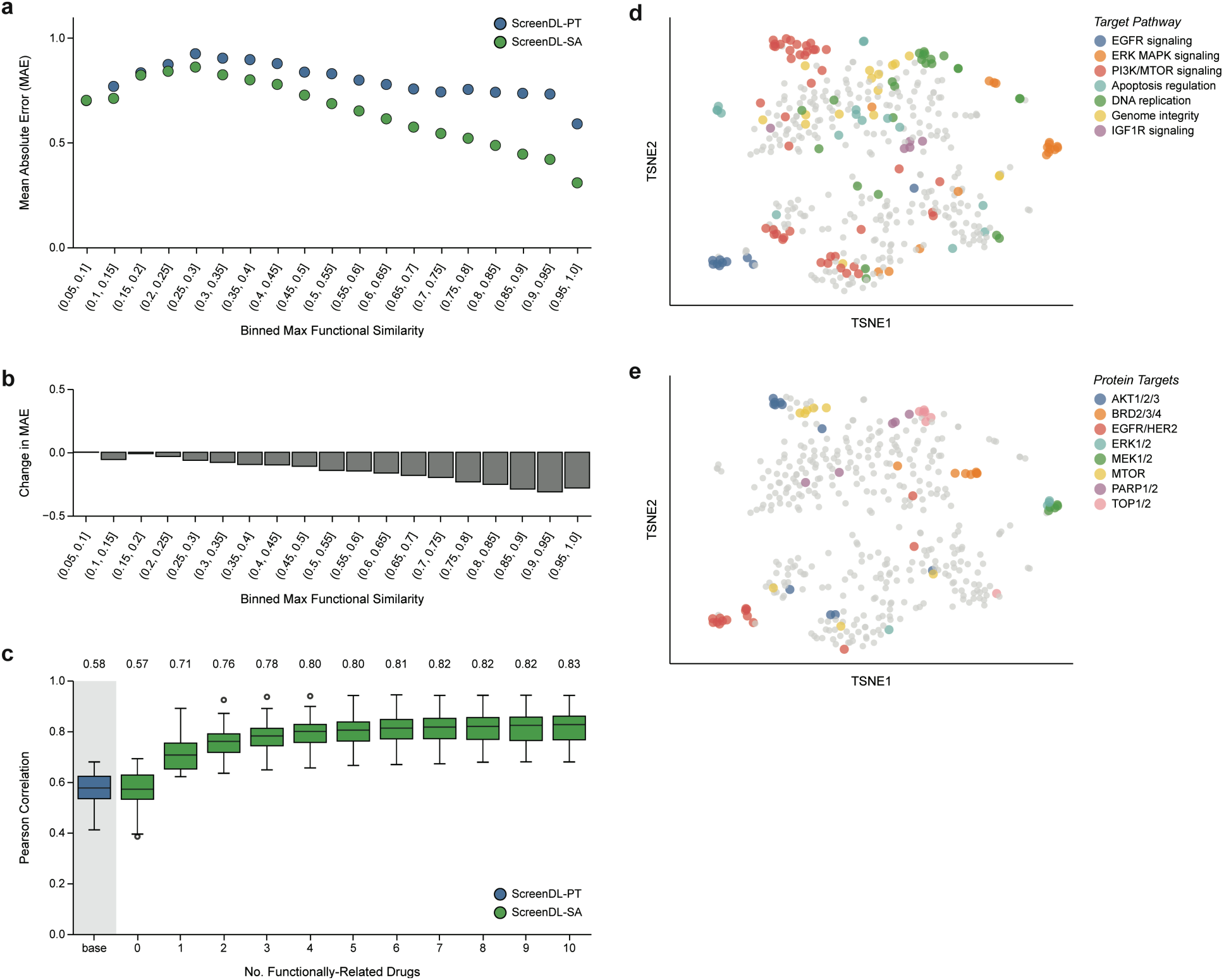
ScreenAhead leverages learned drug-drug functional relationships to improve predictions for unscreened therapies. **a.** Mean absolute error (MAE) of *Z_d_* predictions in cell lines for ScreenDL-PT and ScreenDL-SA. Each cell line-drug pair was assigned to a bin according to the maximum functional similarity with the 20 drugs used for ScreenAhead tumor-specific fine-tuning in the corresponding cell line (see Supplementary Text 3). After assigning each response to a bin (x-axis), the MAE of *Z_d_* predictions for ScreenDL-PT and ScreenDL-SA was computed for each bin (y-axis). ScreenAhead provided outsized performance gains in cell line-drug pairs for which the drug had a higher maximum functional similarity with drugs in the ScreenAhead drug set. **b**. Change in MAE after ScreenAhead (ScreenDL-SA vs ScreenDL-PT) for each bin. **c.** Pearson correlation between observed and predicted *Z_d_* responses for 20 drugs (5-fluorouracil, leflunomide, epirubicin, piperlongumine, vinblastine, oxaliplatin, docetaxel, gemcitabine, cytarabine, cisplatin, alisertib, afatinib, erlotinib, dabrafenib, alpelisib, trametinib, olaparib, nilotinib, fulvestrant, and irinotecan) in cell lines when including an increasing number of functionally related therapies in ScreenAhead tumor-specific fine-tuning. For each interested drug, we compared the performance of ScreenDL-PT at baseline to that achieved by ScreenDL-SA when including an increasing number of functionally related therapies in the ScreenAhead drug set. Performance significantly improved with the inclusion of just one functionally related therapy. Additional improvement was observed upon inclusion of each additional functionally related agent. **d,e.** tSNE plots of ScreenDL-PT’s drug subnetwork embeddings colored by annotations of drug biological mechanism (**d**) or protein targets (**e**).

**Extended Data Fig. 7:**
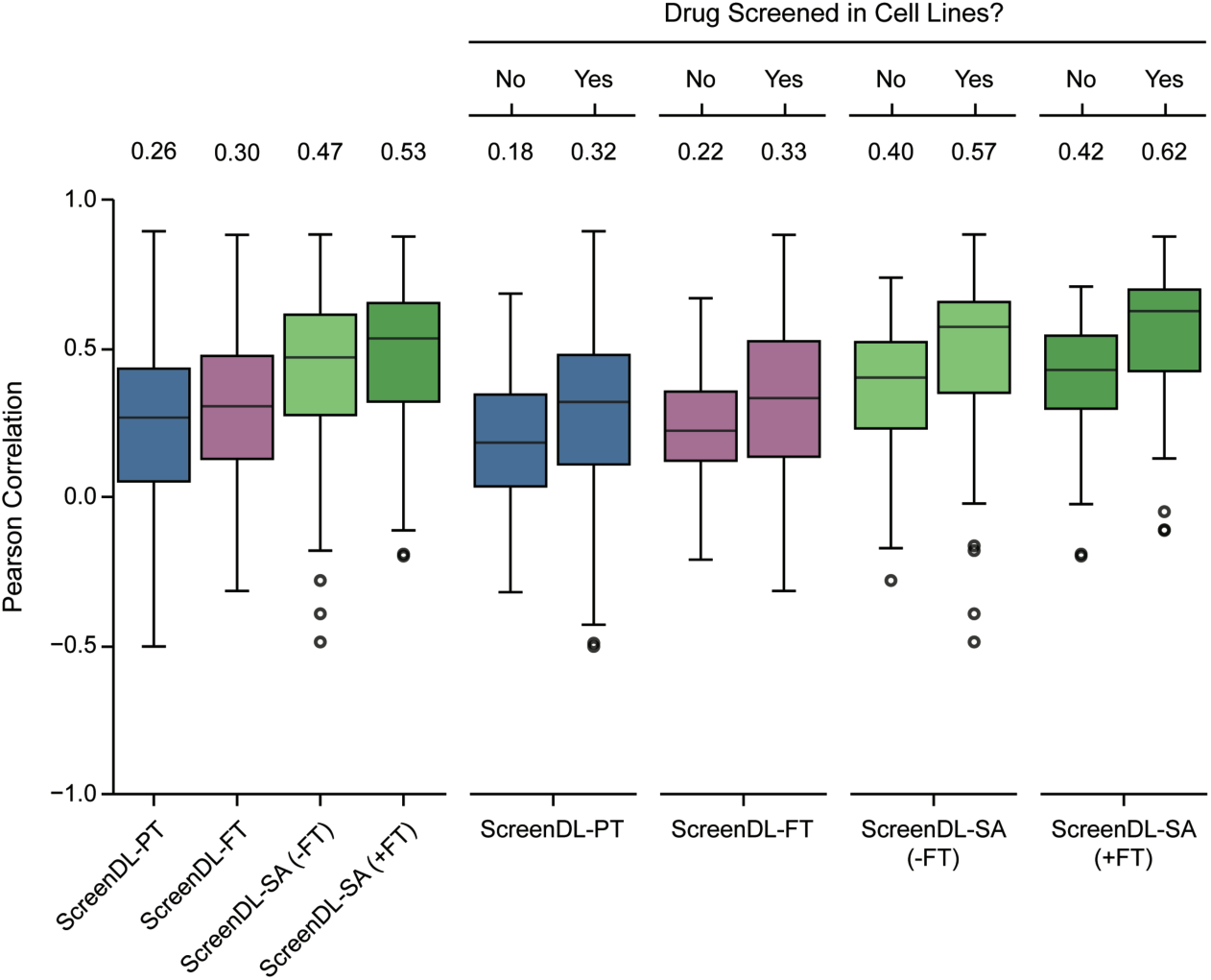
The combination of domain-specific fine-tuning and ScreenAhead tumor-specific fine-tuning is necessary for optimal performance. Drug-level performance of ScreenDL-SA with (+FT) and without (- FT) prior domain-specific fine-tuning. Box plots represent the distribution of Pearson correlations between observed and predicted response per drug. Median values for each model are denoted in the upper margin. Performance is shown across all drugs (left) and stratified according to whether or not the tested drugs were screened in cell lines and thus included in cell line pretraining (right). The performance of ScreenDL-PT and ScreenDL-FT are shown for reference.

## Supplementary Information

### Supplementary Text 1. Evaluation of ScreenAhead under different prediction scenarios

The ScreenAhead approach relies on input of patient-specific data. However, both biological and clinical constraints, including limited biomaterial from tumor sampling and poor organoid growth, may limit the number of therapies that can be effectively screened before treatment decisions are made. Given these potential restrictions on the scope of pre-treatment drug screening, we explored the efficacy of our ScreenAhead approach while incrementally expanding the number of drugs in the partial screening set for each cell line. With as few as five drugs screened, we observed a significant improvement in performance, with ScreenDL-SA achieving a median PCC per drug of 0.59 compared to 0.48 for ScreenDL-PT (p < 0.0005, two-sided Wilcoxon signed-rank test) (Extended Data Fig. 4a). Furthermore, we observed continued improvements in prediction accuracy as an increasing number of drugs were added to the partial screening set for each cell line (Extended Data Fig. 4a).

Given our finding that ScreenAhead enhances model performance in part by utilizing cross-information from biologically related drugs, we reasoned that an informed approach to drug selection would enable ScreenDL-SA to better leverage the limited partial screening data available from functional studies in patient-derived tumor models. Accordingly, we evaluated three informed drug selection algorithms designed to maximize coverage of the drug response space and ensure systematic exploration of potential therapeutic responses: agglomerative clustering, metadata-based selection using target pathway annotations, and Principal Feature Analysis (PFA)^47^. We conducted comparative analyses of drug response predictions with ScreenDL-SA under two scenarios: random drug selection, and informed selection using each of these three algorithms. Our results illustrate that drug selection using PFA consistently outperforms random selection, with PFA achieving superior performance for 68% of drugs when including 20 drugs in partial screening (Extended Data Fig. 4c). Furthermore, a pairwise analysis of drug-level performance revealed that PFA significantly outperforms drug selection using either hierarchical clustering or target pathway annotations, particularly as the size of the pre-screening drug set increases (Extended Data Fig. 4d).

### Supplementary Text 2. ScreenAhead leverages knowledge of GDS to calibrate response predictions

To investigate the role of global drug sensitivity (GDS) in ScreenAhead, we stratified the performance of ScreenDL-PT and ScreenDL-SA according to the expected performance of a GDS-only model. Specifically, for each cell line *c*, we defined global drug sensitivity *GDS*_*c*_ as the cell line’s mean observed *Z*_*d*_ across drugs and generated a GDS-only model *f*(*c*, *d*) = *GDS*_*c*_that always returns a given cell line’s GDS as the predicted response regardless of drug features. We then grouped cell line-drug pairs by binning the absolute error between observed and predicted *Z*_*d*_ values for this GDS-only model and computed the mean absolute error (MAE) of predictions within each bin for ScreenDL-PT and ScreenDL-SA. Extended Data Fig. 5d reveals that ScreenAhead improves performance, even for cell line-drug pairs for which GDS was not highly predictive. To further characterize the role of GDS in ScreenAhead, we performed tumor-specific fine-tuning for each cell line in our harmonized pharmaco-omic dataset under two scenarios: (1) using the cell line’s true *Z*_*d*_ responses for 20 drugs; or (2) replacing the true *Z*_*d*_ responses with the cell line’s mean *Z*_*d*_ across these 20 drugs. In the later scenario, the use of mean-filled *Z*_*d*_ responses removes drug-specific information encoded in a cell line’s partial screening data and instead provides only an estimate of GDS during tumor-specific fine-tuning. This analysis revealed that, while incorporating knowledge of GDS through ScreenAhead improved performance, GDS alone did not account for the full performance gain seen with ScreenDL-SA (Extended Data Fig. 5c).

### Supplementary Text 3. ScreenAhead exploits cross-information from functionally related therapies

To determine if ScreenAhead tumor-specific fine-tuning enhanced predictions for unscreened drugs (i.e., those drugs not included in ScreenAhead for a given tumor) by leveraging learned drug-drug functional similarities to borrow information from pre-screened therapies, we stratified cell line performance based on the maximum functional similarity of a given unscreened drug with the 20 drugs selected for ScreenAhead tumor-specific fine-tuning for the corresponding cell line. Concretely, given the sets of cell lines *C*, drugs *D*, and observed responses *R* = {*r*(*c*, *d*) ∣ *c* ∈ *C*, *d* ∈ *D*}, each response *r*(*c*, *d*) was assigned to a bin according to the maximum functional similarity *F*_*max*_(*c*, *d*) = *max*_*s_i_*∈*S_c_*_{*F*(*d*, *s*_*i*_)} between the drug *d* and another drug *s*_*i*_ belonging to the set of ScreenAhead drugs *S*_*c*_ = {*s*_1_, *s*_2_, . . ., *s*_20_} for the cell line *c*. Functional similarity *F*(*d*, *s*_*i*_) was defined as the Pearson correlation coefficient (PCC) between the observed cell line response profiles for *d* and *s*_*i*_. After assigning each response to a bin, we computed the mean absolute error (MAE) of *Z*_*d*_ predictions for ScreenDL-PT and ScreenDL-SA separately for each bin. Performance gains from ScreenAhead were strongly correlated with *F*_*max*_(*c*, *d*) (PCC = - 0.98 between the observed change in MAE after ScreenAhead and bin rank, p = 9.85 x 10^-14^; Extended Data Fig. 6a,b).

To further characterize the positive transfer of information between functionally related therapies during ScreenAhead, we selected 20 drugs for a focused analysis: 5-fluorouracil, leflunomide, epirubicin, piperlongumine, vinblastine, oxaliplatin, docetaxel, gemcitabine, cytarabine, cisplatin, alisertib, afatinib, erlotinib, dabrafenib, alpelisib, trametinib, olaparib, nilotinib, fulvestrant, and irinotecan. For each interested drug *d*, we generated an initial pre-screening set containing the 20 least functionally similar drugs in our cell line dataset by ranking drugs according to the PCC between their observed cell line responses and those of the interested drug *d*. We then performed ScreenAhead tumor-specific fine-tuning with this initial drug set and with 10 modified pre-screening drug sets generated by replacing an increasing number of drugs with drug *d*’s most functionally related therapies. Relative to ScreenDL-PT, ScreenAhead tumor-specific fine-tuning did not improve performance for the interested drugs when no functionally similar therapies were included in the ScreenAhead (p = 0.67, Wilcoxon signed-rank test). In comparison, we observed significant improvement when just one functionally related therapy was included (p = 1.91 x 10^-6^, Wilcoxon signed-rank test; Extended Data Fig. 6c). Further, we observed additional performance gains with the inclusion of each successive functionally related therapy (Extended Data Fig. 6c), illustrating ScreenDL’s ability to exploit cross-information from functionally related drugs during ScreenAhead tumor-specific fine-tuning.

### Supplementary Text 4. Calculation of PDXO dose-response metrics

PDXO drug response was quantified by calculating IC50 and growth rate (GR)-adjusted area over the curve (GRaoc) values for each PDXO-drug pair^21^. IC50, GR, and subsequent GRaoc values were calculated using the GRmetrics R package^57^. While natural log transformed IC50 values were used in the development of ScreenDL to facilitate the harmonization of PDXO and cell line drug response data, the calculation of GRaoc values for cBioPortal involved several modifications to improve the stability of GR calculations and reduce bias in GRaoc values. Specifically, GRaoc was quantified by measuring drug-induced normalized growth rate inhibition (GR) for each drug concentration and integrating over the concentration curve to provide a GRaoc value for each drug as described by Hafner and colleagues^66,67^. Normalization values for GR calculations were established using dimethyl sulfoxide (DMSO) dosed control wells.

Due to the mathematics involved, GR values become unstable when control cell count fold change is close to 1. To limit this instability, we developed two criteria that must be met for inclusion of a plate in the final drug screening dataset. First, the average fold change in control wells on a plate must be greater than 1.5. Second, the overlap in kernel density estimates between baseline count values and control count values for a plate must be less than 30%. The cutoff for these filters was established graphically by looking for the inflection point where GR values went from wide variation to clustering closely around 1 for individual DMSO wells using average plate DMSO count as the control.

GR values are expected to be between -1 and 1. GR should be 1 if a drug at a given concentration has no effect on the growth rate of cells. If, by random chance, the number of cells in a treated well with no effect is greater than the control well, then GR can be greater than 1. In some cases, GR can be much greater than 1 and has an undue influence on GRaoc. The function used to calculate GRaoc in the GRmetrics package has an option to cap GR values at 1. We found that capping GR values at 1 biases GRaoc values to be high; additionally, PDXO models with low control fold change have more bias than PDXO models with higher control fold change. The 95th percentile of individual DMSO wells using average plate DMSO count as the control was 1.6. We capped GR at 1.6 which results in far less bias in GRaoc values and more uniform bias across control fold change values.

## Supplementary Figures

**Supplementary Fig. 1:**
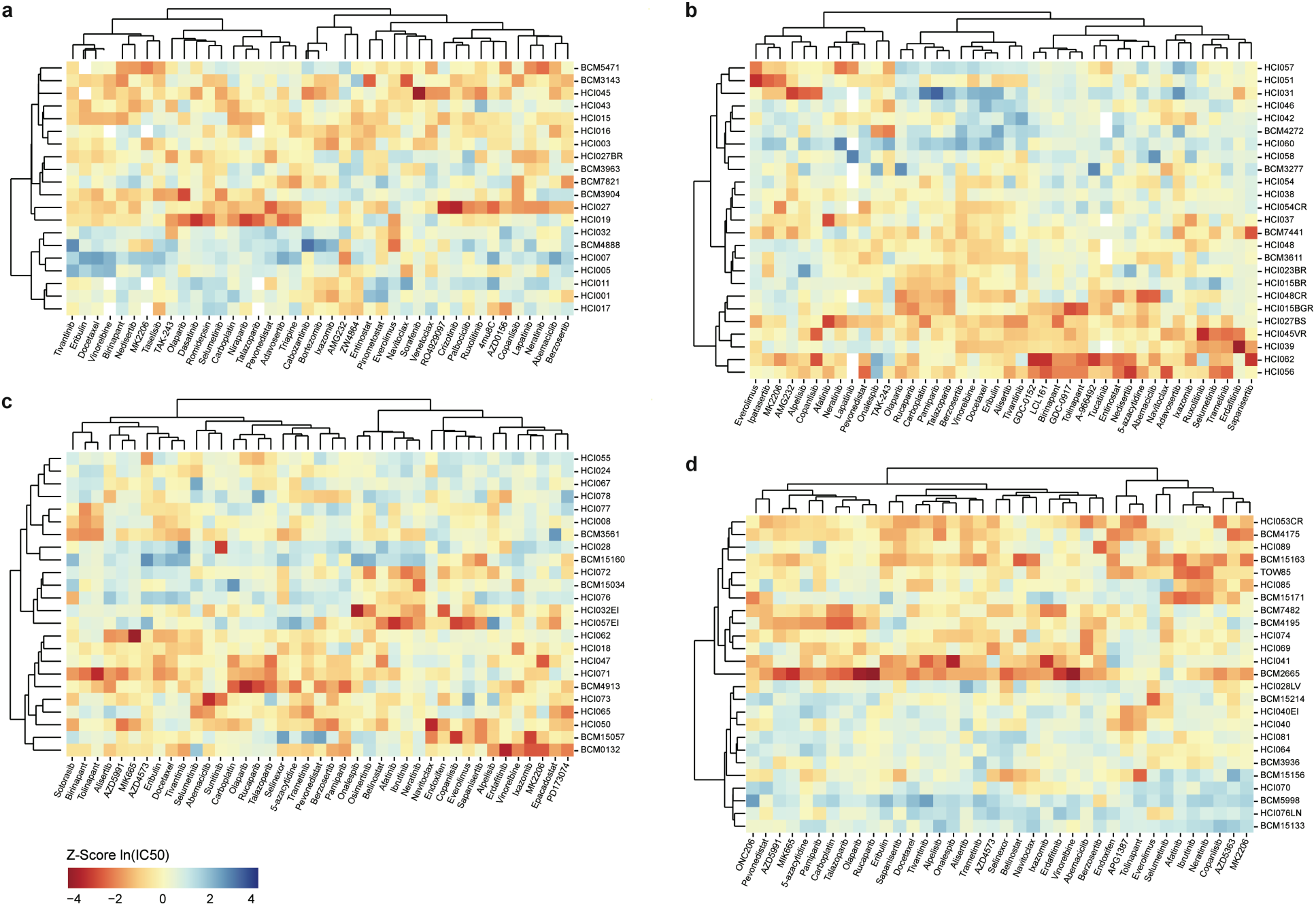
Raw PDxO screening data. a-d. Unsupervised clustering of the PDXOs and drugs screened in each of four screens A-D (**a-d;** see Methods). Color indicates the *z*-score normalized ln(IC50) value for a given PDXO-drug pair (darker red indicates cytotoxicity and darker blue indicates growth).

## References

1. Suehnholz, S. P. et al. Quantifying the Expanding Landscape of Clinical Actionability for Patients with Cancer. Cancer Discov. 14, 49–65 (2024).

2. Zehir, A. et al. Mutational landscape of metastatic cancer revealed from prospective clinical sequencing of 10,000 patients. Nat. Med. 23, 703–713 (2017).

3. Haslam, A., Kim, M. S. & Prasad, V. Updated estimates of eligibility for and response to genome-targeted oncology drugs among US cancer patients, 2006-2020. Ann. Oncol. 32, 926–932 (2021).

4. Marquart, J., Chen, E. Y. & Prasad, V. Estimation of the Percentage of US Patients With Cancer Who Benefit From Genome-Driven Oncology. JAMA Oncol 4, 1093–1098 (2018).

5. Flaherty, K. T. et al. Molecular Landscape and Actionable Alterations in a Genomically Guided Cancer Clinical Trial: National Cancer Institute Molecular Analysis for Therapy Choice (NCI-MATCH). J. Clin. Oncol. 38, 3883–3894 (2020).

6. Schwaederle, M. et al. Association of Biomarker-Based Treatment Strategies With Response Rates and Progression-Free Survival in Refractory Malignant Neoplasms: A Meta-analysis. JAMA Oncol 2, 1452–1459 (2016).

7. Yancovitz, M. et al. Intra- and inter-tumor heterogeneity of BRAF(V600E))mutations in primary and metastatic melanoma. PLoS One 7, e29336 (2012).

8. Fisher, R., Pusztai, L. & Swanton, C. Cancer heterogeneity: implications for targeted therapeutics. Br. J. Cancer 108, 479–485 (2013).

9. Ganesh, K. & Massagué, J. Targeting metastatic cancer. Nat. Med. 27, 34–44 (2021).

10. Le Tourneau, C. et al. Molecularly targeted therapy based on tumour molecular profiling versus conventional therapy for advanced cancer (SHIVA): a multicentre, open-label, proof-of-concept, randomised, controlled phase 2 trial. Lancet Oncol. 16, 1324–1334 (2015).

11. Pezo, R. C. et al. Impact of multi-gene mutational profiling on clinical trial outcomes in metastatic breast cancer. Breast Cancer Res. Treat. 168, 159–168 (2018).

12. Ding, M. Q., Chen, L., Cooper, G. F., Young, J. D. & Lu, X. Precision Oncology beyond Targeted Therapy: Combining Omics Data with Machine Learning Matches the Majority of Cancer Cells to Effective Therapeutics. Mol. Cancer Res. 16, 269–278 (2018).

13. Adashek, J. J., Goloubev, A., Kato, S. & Kurzrock, R. Missing the target in cancer therapy. *Nat*. Cancer 2, 369–371 (2021).

14. Partin, A. et al. Deep learning methods for drug response prediction in cancer: Predominant and emerging trends. Front. Med. 10, 1086097 (2023).

15. Adam, G. et al. Machine learning approaches to drug response prediction: challenges and recent progress. NPJ Precis Oncol 4, 19 (2020).

16. Ovchinnikova, K., Born, J., Chouvardas, P., Rapsomaniki, M. & Kruithof-de Julio, M. Overcoming limitations in current measures of drug response may enable AI-driven precision oncology. NPJ Precis Oncol 8, 95 (2024).

17. Wilding, J. L. & Bodmer, W. F. Cancer cell lines for drug discovery and development. Cancer Res. 74, 2377–2384 (2014).

18. Abbas, Z. N., Al-Saffar, A. Z., Jasim, S. M. & Sulaiman, G. M. Comparative analysis between 2D and 3D colorectal cancer culture models for insights into cellular morphological and transcriptomic variations. Sci. Rep. 13, 18380 (2023).

19. Hidalgo, M. et al. Patient-derived xenograft models: an emerging platform for translational cancer research. Cancer Discov. 4, 998–1013 (2014).

20. Gao, H. et al. High-throughput screening using patient-derived tumor xenografts to predict clinical trial drug response. Nat. Med. 21, 1318–1325 (2015).

21. Guillen, K. P. et al. A human breast cancer-derived xenograft and organoid platform for drug discovery and precision oncology. Nat Cancer 3, 232–250 (2022).

22. Wensink, G. E. et al. Patient-derived organoids as a predictive biomarker for treatment response in cancer patients. NPJ Precis Oncol 5, 30 (2021).

23. Lee, J.-K. et al. Pharmacogenomic landscape of patient-derived tumor cells informs precision oncology therapy. Nat. Genet. 50, 1399–1411 (2018).

24. Bruna, A. et al. A Biobank of Breast Cancer Explants with Preserved Intra-tumor Heterogeneity to Screen Anticancer Compounds. Cell 167, 260–274.e22 (2016).

25. Pauli, C. et al. Personalized In Vitro and In Vivo Cancer Models to Guide Precision Medicine. Cancer Discov. 7, 462–477 (2017).

26. Wang, E., Xiang, K., Zhang, Y. & Wang, X.-F. Patient-derived organoids (PDOs) and PDO-derived xenografts (PDOXs): New opportunities in establishing faithful pre-clinical cancer models. Journal of the National Cancer Center 2, 263–276 (2022).

27. Margossian, A. et al. Clinical and genomic correlation of a CLIA certified organoid based functional test in breast cancer patients. Cancer Res. 81, PS4–01 (2021).

28. Margossian, A. et al. Predictive value of a CLIA-approved organoid based drug sensitivity test. J. Clin. Oncol. 38, 3630–3630 (2020).

29. Letai, A., Bhola, P. & Welm, A. L. Functional precision oncology: Testing tumors with drugs to identify vulnerabilities and novel combinations. Cancer Cell 40, 26–35 (2022).

30. Turner, R. M., Park, B. K. & Pirmohamed, M. Parsing interindividual drug variability: an emerging role for systems pharmacology. Wiley Interdiscip. Rev. Syst. Biol. Med. 7, 221– 241 (2015).

31. Rogers, D. & Hahn, M. Extended-connectivity fingerprints. J. Chem. Inf. Model. 50, 742– 754 (2010).

32. Liberzon, A. et al. The Molecular Signatures Database (MSigDB) hallmark gene set collection. Cell Syst 1, 417–425 (2015).

33. Hostallero, D. E., Li, Y. & Emad, A. Looking at the BiG picture: incorporating bipartite graphs in drug response prediction. Bioinformatics 38, 3609–3620 (2022).

34. Yang, W. et al. Genomics of Drug Sensitivity in Cancer (GDSC): a resource for therapeutic biomarker discovery in cancer cells. Nucleic Acids Res. 41, D955–61 (2013).

35. van der Meer, D. et al. Cell Model Passports—a hub for clinical, genetic and functional datasets of preclinical cancer models. Nucleic Acids Res. 47, D923–D929 (2018).

36. Liu, Q., Hu, Z., Jiang, R. & Zhou, M. DeepCDR: a hybrid graph convolutional network for predicting cancer drug response. Bioinformatics 36, i911–i918 (2020).

37. Ma, T. et al. DualGCN: a dual graph convolutional network model to predict cancer drug response. BMC Bioinformatics 23, 129 (2022).

38. Jin, I. & Nam, H. HiDRA: Hierarchical Network for Drug Response Prediction with Attention. J. Chem. Inf. Model. 61, 3858–3867 (2021).

39. Scherer, S. D. et al. Breast cancer PDxO cultures for drug discovery and functional precision oncology. STAR Protoc. 4, 102402 (2023).

40. Cerami, E. et al. The cBio cancer genomics portal: an open platform for exploring multidimensional cancer genomics data. Cancer Discov. 2, 401–404 (2012).

41. Gao, J. et al. Integrative analysis of complex cancer genomics and clinical profiles using the cBioPortal. Sci. Signal. 6, l1 (2013).

42. de Bruijn, I. et al. Analysis and visualization of longitudinal genomic and clinical data from the AACR Project GENIE Biopharma Collaborative in cBioPortal. Cancer Res. 83, 3861– 3867 (2023).

43. Turner Nicholas C., et al. Capivasertib in Hormone Receptor–Positive Advanced Breast Cancer. N. Engl. J. Med. 388, 2058–2070 (2023).

44. Vaklavas, C. et al. TOWARDS study: Patient-derived xenograft engraftment predicts poor survival in patients with newly diagnosed triple-negative breast cancer. *JCO Precis*. Oncol. 8, e2300724 (2024).

45. Geeleher, P., Cox, N. J. & Huang, R. S. Cancer biomarker discovery is improved by accounting for variability in general levels of drug sensitivity in pre-clinical models. Genome Biol. 17, 190 (2016).

46. White, B. S. et al. Bayesian multi-source regression and monocyte-associated gene expression predict BCL-2 inhibitor resistance in acute myeloid leukemia. NPJ Precis Oncol 5, 71 (2021).

47. Lu, Y., Cohen, I., Zhou, X. S. & Tian, Q. Feature selection using principal feature analysis. in Proceedings of the 15th ACM international conference on Multimedia 301–304 (Association for Computing Machinery, New York, NY, USA, 2007).

48. Bose, S. et al. A path to translation: How 3D patient tumor avatars enable next generation precision oncology. Cancer Cell 40, 1448–1453 (2022).

49. Cocco, S. et al. Biomarkers in Triple-Negative Breast Cancer: State-of-the-Art and Future Perspectives. Int. J. Mol. Sci. 21, (2020).

50. Meric-Bernstam, F. et al. Assessment of patient-derived xenograft growth and antitumor activity: The NCI PDXNet consensus recommendations. Mol. Cancer Ther. 23, 924–938 (2024).

51. Narasimhan, V. et al. Medium-throughput drug screening of patient-derived organoids from colorectal peritoneal metastases to direct personalized therapy. Clin. Cancer Res. 26, 3662–3670 (2020).

52. Tiriac, H. et al. Organoid profiling identifies common responders to chemotherapy in pancreatic cancer. Cancer Discov. 8, 1112–1129 (2018).

53. He, X. et al. Patient-derived organoids as a platform for drug screening in metastatic colorectal cancer. Front Bioeng Biotechnol 11, 1190637 (2023).

54. Broutier, L. et al. Human primary liver cancer-derived organoid cultures for disease modeling and drug screening. Nat. Med. 23, 1424–1435 (2017).

55. Kim, S. et al. PubChem 2023 update. Nucleic Acids Res. 51, D1373–D1380 (2023).

56. Behdenna, A. et al. pyComBat, a Python tool for batch effects correction in high-throughput molecular data using empirical Bayes methods. BMC Bioinformatics 24, 459 (2023).

57. Clark, N. A. et al. GRcalculator: an online tool for calculating and mining dose-response data. BMC Cancer 17, 698 (2017).

58. Partin, A. et al. Learning curves for drug response prediction in cancer cell lines. BMC Bioinformatics 22, 252 (2021).

59. Jang, I. S., Neto, E. C., Guinney, J., Friend, S. H. & Margolin, A. A. Systematic assessment of analytical methods for drug sensitivity prediction from cancer cell line data. Pac. Symp. Biocomput. 63–74 (2014).

60. Costello, J. C. et al. A community effort to assess and improve drug sensitivity prediction algorithms. Nat. Biotechnol. 32, 1202–1212 (2014).

61. DeRose, Y. S. et al. Patient-derived models of human breast cancer: protocols for in vitro and in vivo applications in tumor biology and translational medicine. Curr. Protoc. Pharmacol. Chapter 14, Unit14.23 (2013).

62. DeRose, Y. S. et al. Tumor grafts derived from women with breast cancer authentically reflect tumor pathology, growth, metastasis and disease outcomes. Nat. Med. 17, 1514– 1520 (2011).

63. Nishino, M., Jagannathan, J. P., Ramaiya, N. H. & Van den Abbeele, A. D. Revised RECIST guideline version 1.1: What oncologists want to know and what radiologists need to know. AJR Am. J. Roentgenol. 195, 281–289 (2010).

64. McLaren, W. et al. The Ensembl Variant Effect Predictor. Genome Biol. 17, 122 (2016).

65. Evrard, Y. A. et al. Systematic establishment of robustness and standards in patient-derived xenograft experiments and analysis. Cancer Res. 80, 2286–2297 (2020).

66. Hafner, M., Niepel, M., Chung, M. & Sorger, P. K. Growth rate inhibition metrics correct for confounders in measuring sensitivity to cancer drugs. Nat. Methods 13, 521–527 (2016).

67. Hafner, M., Niepel, M. & Sorger, P. K. Alternative drug sensitivity metrics improve preclinical cancer pharmacogenomics. Nat. Biotechnol. 35, 500–502 (2017).

