## Supplementary Table S1. Summary of HCI breast cancer PDXO and PDX lines with clinical annotations for "A precision oncology-focused deep learning framework for personalized selection of cancer therapy"

Table 1. Summary of current HCI breast cancer PDXO and PDX lines, along with sublines.

| | Patient tumor information | | | | | | PDX information | | | | PDXO information | | | |
|---|---|---|---|---|---|---|---|---|---|---|---|---|---|---|
| | Model ID | Collection site | Tx naïve or pretreated | ER | PR | HER2 | ER | PR | HER2 | Estrogen Dependent | PDXO information | Doubling time | Passage interval | PDXO drug screened |
| 1 | HCI-001 | 1° breast | Pretreated | neg | neg | neg | neg | neg | neg | n/a | Yes | 3–5 days | 12–15 days | Yes |
| 2 | HCI-002 | 1° breast | Naïve | neg | neg | neg | neg | neg | neg | n/a | Yes | 2–4 days | 10–12 days | Yes |
| 3 | HCI-003 | 1° breast | Naïve | pos | pos | neg | pos | pos | neg | Yes; ESR1 WT | Yes | 4–7 days | 15–20 days | Yes |
| 4 | HCI-003FR* | 1° breast | Naïve | pos | pos | neg | pos | pos | neg | Yes; ESR1 WT | not attempted | | | |
| 5 | HCI-004 | 1° breast | Naïve | neg | neg | neg | neg | neg | neg | n/a | Failed (no organoid formation) | | | |
| 6 | HCI-005 | Pleural Effusion | Pretreated | pos | pos | primary tumor was pos and amplified; PE not tested | pos | pos | pos, mixed; not amp | Partial; E2 enhanced; ESR1 L536P | Yes | 7–10 days | 20–25 days | Yes |
| 7 | HCI-006 | Pleural Effusion | Pretreated | pos | pos | primary tumor was pos and amplified; PE not tested | pos | pos | nd | nd; ESR1 L536P | Failed (no organoid formation) | | | |
| 8 | HCI-007 | Pleural Effusion | Pretreated | pos | pos | primary tumor was pos and amplified; PE not tested | pos | pos | pos, mixed; not amp | nd; ESR1 L536P | Yes | 7–10 days | 20–25 days | Yes |
| 9 | HCI-008 | Pleural Effusion | Pretreated | neg | neg | pos | neg | neg | pos, mixed; not amp | n/a | Yes | 4–7 days | 15–20 days | Yes |
| 10 | HCI-009 | Ascites | Pretreated | neg | neg | neg | neg | neg | neg | n/a | Failed (no organoid formation) | | | |
| 11 | HCI-010 | Pleural Effusion | Pretreated | neg | neg | neg | neg | borderline | neg | n/a | Yes | 4–7 days | 15–20 days | Yes |
| 12 | HCI-011 | Pleural Effusion | Pretreated | pos | pos | neg | pos | pos | neg | Partial; E2 enhanced; ESR1 WT | Yes | 7–10 days | 20–25 days | Yes |
| 13 | HCI-011FR* | Pleural Effusion | Pretreated | pos | pos | neg | pos | pos | neg | Partial; E2 enhanced; ESR1 WT | not attempted | | | |
| 14 | HCI-012 | Pleural Effusion | Pretreated | neg | neg | primary tumor was pos and amplified; PE not tested | neg | neg | pos, mixed | n/a | Yes | 4–7 days | 15–20 days | Yes |
| 15 | HCI-013 | Pleural Effusion | Pretreated | pos | pos | neg | pos | pos | neg | Partial; E2 enhanced; ESR1 [illegible] | Failed (no organoid formation) | | | |
| 16 | HCI-013EI | Pleural Effusion | Pretreated | pos | pos | neg | pos | pos | neg | Partial; E2 enhanced; ESR1 [illegible] | not attempted | | | |
| 17 | HCI-014 | Pleural Effusion | Pretreated | neg | neg | neg | neg | neg | neg | n/a | Failed (no organoid formation) | | | |
| 18 | HCI-015 | Brain metastasis | Pretreated | neg | neg | neg | neg | neg | neg | n/a | Yes | 4–7 days | 15–20 days | Yes |
| 19 | HCI-015BR | Brain metastasis | Pretreated | neg | neg | neg | neg | neg | neg | n/a | Yes | 4–7 days | 15–20 days | Yes |
| 20 | HCI-015BGR | Brain metastasis | Pretreated | neg | neg | neg | neg | neg | neg | n/a | Yes | 3–5 days | 12–15 days | Yes |
| 21 | HCI-016 | 1° breast | Unknown | neg | neg | neg | neg | neg | neg | n/a | Yes | 4–7 days | 15–20 days | Yes |
| 22 | HCI-017 | 1° breast | Naïve | pos | pos | neg | pos | pos | neg | Yes; ESR1 WT | Yes | 4–7 days | 15–20 days | Yes |
| 23 | HCI-017B | Bone metastasis | Pretreated | pos | pos | neg | pos | nd | neg | nd | not attempted | | | |
| 24 | HCI-018 | Brain metastasis | Pretreated | pos | neg | neg | pos | neg | neg | Yes; ESR1 WT (4-9% Y537S) | Yes | 10–15 days | 25–30 days | Yes |
| 25 | HCI-018BRN | Brain metastasis | Pretreated | pos | neg | neg | pos | neg | neg | Yes; ESR1 WT (4-9% Y537S) | Yes | 10–15 days | 25–30 days | no |
| 26 | HCI-019 | 1° breast | Naïve | neg (1%) | neg (1%) | neg | neg | neg | neg | n/a | Yes | 7–10 days | 20–25 days | Yes |
| 27 | HCI-023 | 1° breast | Naïve | neg | neg | neg | neg | neg | neg | n/a | Yes | 7–10 days | 20–25 days | Yes |
| 28 | HCI-023BR | 1° breast | Naïve | neg | neg | neg | neg | neg | neg | n/a | Yes | 4–7 days | 15–20 days | Yes |
| 29 | HCI-024 | Skin metastasis | Pretreated | neg | neg | neg | neg | neg | neg | n/a | Yes | 4–7 days | 15–20 days | Yes |
| 30 | HCI-025 | Skin metastasis | Pretreated | neg | neg | neg | neg | neg | neg | n/a | Yes | 4–7 days | 15–20 days | Yes |
| 31 | HCI-026 | Spine metastasis | Pretreated | pos | neg (3%) | neg | pos | neg | neg | nd; ESR1 WT | Failed (no organoid formation) | | | |
| 32 | HCI-027 | 1° breast | Pretreated | neg | neg | neg | neg | neg | neg | n/a | Yes | 3–5 days | 12–15 days | Yes |
| 33 | HCI-027BR | 1° breast | Pretreated | neg | neg | neg | neg | neg | neg | n/a | Yes | 4–7 days | 15–20 days | Yes |
| 34 | HCI-027BS | 1° breast | Pretreated | neg | neg | neg | neg | neg | neg | n/a | Yes | 7–10 days | 20–25 days | Yes |
| 35 | HCI-028 | Pleural Effusion | Pretreated | neg | neg | neg | neg | neg | neg | n/a | Yes | 7–10 days | 20–25 days | Yes |
| 36 | HCI-028LV | Pleural Effusion | Pretreated | neg | neg | neg | neg | neg | neg | n/a | Yes | 10–15 days | 25–30 days | Yes |
| 37 | HCI-030 | recurr. breast | Pretreated | neg | neg | neg | neg | neg | neg | n/a | Failed (no organoid formation) | | | |
| 38 | HCI-031 | Pleural Effusion | Pretreated | neg | neg | neg | neg | neg | neg | n/a | Yes | 15–19 days | 25–30 days | Yes |
| 39 | HCI-031OV | Pleural Effusion | Pretreated | neg | neg | neg | neg | neg | neg | n/a | Yes | 12-14 days | 30-35 | no |
| 40 | HCI-032 | recurr. breast | Pretreated | pos | pos | pos | pos | pos | pos | No; ESR1 WT (1-3% Y537S) | Yes | 4–7 days | 15–20 days | Yes |
| 41 | HCI-032EI | recurr. breast | Pretreated | pos | pos | pos | pos | pos | pos | No; ESR1 WT (1-3% Y537S) | Yes | 7–10 days | 20–25 days | Yes |
| 42 | HCI-033 | Lung metastasis | Pretreated | neg | neg | neg | neg | neg | neg | n/a | Failed (no organoid formation) | | | |
| 43 | HCI-034 | recurr. breast | Pretreated | neg | neg | neg | neg | neg | neg | n/a | Failed (no organoid formation) | | | |
| 44 | HCI-036 | recurr. breast phyllodes | Naïve | neg | neg | neg | neg | neg | neg | n/a | Failed (no organoid formation) | | | |
| 45 | HCI-037 | recurr. breast | Pretreated | neg | neg | neg | neg | neg | neg | n/a | Yes | 4–7 days | 15–20 days | Yes |
| 46 | HCI-038 | Bone metastasis | Pretreated | neg | neg | neg | neg | neg | neg | n/a | Yes | 4–7 days | 15–20 days | Yes |
| 47 | HCI-039 | Pleural effusion | Pretreated | neg | neg | neg | neg | neg | neg | n/a | Yes | 4–7 days | 15–20 days | Yes |
| 48 | HCI-040 | Bone metastasis | Pretreated | pos | pos | pos | pos | pos | pos | No; ESR1 WT | Yes | 7–10 days | 20–25 days | Yes |
| 49 | HCI-040EI | Bone metastasis | Pretreated | pos | pos | pos | pos | pos | pos | No; ESR1 WT | Yes | 4–7 days | 15–20 days | Yes |
| 50 | HCI-041 | 1° breast | Naïve | neg | neg | neg | neg | neg | neg | n/a | Yes | 7–10 days | 20–25 days | Yes |
| 51 | HCI-042 | 1° breast | Naïve | neg | neg (1%) | pos | neg | neg | neg | n/a | Yes | 7–10 days | 20–25 days | Yes |
| 52 | HCI-043 | 1° breast | Naïve | neg | neg | neg | neg | neg | neg | n/a | Yes | 7–10 days | 20–25 days | Yes |
| 53 | HCI-044 | Brain metastasis | Pretreated | pos | pos | neg | pos | pos | neg | No; ESR1 Y537S homo | Failed (only mouse content) | | | |
| 54 | HCI-044EI | Brain metastasis | Pretreated | pos | pos | neg | pos | pos | neg | No; ESR1 Y537S homo | Failed (only mouse content) | | | |
| 55 | HCI-044EILV | Brain metastasis | Pretreated | pos | pos | neg | pos | pos | neg | No; ESR1 Y537S homo | not attempted | | | |
| 56 | HCI-045 | 1° breast | Naïve | neg | neg | neg | neg | neg | neg | n/a | Yes | 4–7 days | 15–20 days | Yes |
| 57 | HCI-045VR | 1° breast | Naïve | neg | neg | neg | neg | neg | neg | n/a | Yes | 4–7 days | 15–20 days | Yes |
| 58 | HCI-046 | 1° breast | Naïve | neg | neg | neg | neg | neg | neg | n/a | Yes | 25–30 days | 30–35 days | Yes |
| 59 | HCI-047 | Ascites | Pretreated | neg | neg | neg | neg | neg | neg | n/a | Yes | 4–7 days | 15–20 days | Yes |
| 60 | HCI-048 | 1° breast | Naïve | pos (10%) | neg | neg | neg | neg | neg | n/a | Yes | 4–7 days | 15–20 days | Yes |
| 61 | HCI-048CR | 1° breast | Naïve | pos (10%) | neg | neg | neg | neg | neg | n/a | Yes | 7–10 days | 20–25 days | Yes |
| 62 | HCI-049 | 1° breast | Naïve | pos | pos | neg | pos | pos | neg | Yes; ESR1 WT | Failed (only mouse content) | | | |
| 63 | HCI-050 | LN metastasis | Pretreated | neg (5%) | neg | neg | neg | neg | neg | n/a | Yes | 7–10 days | 20–25 days | Yes |
| 64 | HCI-051 | Ascites | Pretreated | neg | neg | neg | neg | neg | neg | n/a | Yes | 4–7 days | 15–20 days | Yes |
| 65 | HCI-052 | 1° breast | Naïve | neg | neg | neg | neg | neg | neg | n/a | Failed (no organoid formation) | | | |
| 66 | HCI-053 | 1° breast | Naïve | neg | neg | neg | neg | neg | neg | n/a | Yes | 4–7 days | 15–20 days | Yes |
| 67 | HCI-053CR | 1° breast | Naïve | neg | neg | neg | neg | neg | neg | n/a | Yes | 7–10 days | 20–25 days | Yes |
| 68 | HCI-054 | 1° breast | Pretreated | neg | neg | neg | neg | neg | neg | n/a | Yes | 10–15 days | 25–30 days | Yes |
| 69 | HCI-054CR | 1° breast | Pretreated | neg | neg | neg | neg | neg | neg | n/a | Yes | 7–10 days | 20–25 days | Yes |
| 70 | HCI-055 | 1° breast | Naïve | neg | neg | neg | neg | neg | neg | n/a | yes | 7–10 days | 20–25 days | yes |
| 71 | HCI-056 | 1° breast | Naïve | neg | neg | neg | neg | neg | neg | n/a | yes | 6–7 days | 15–20 days | yes |
| 72 | HCI-057 | Liver metastasis | Pretreated | pos | pos | pos | pos | pos | pos | No; ESR1 WT | yes | 7–14 days | 20–30 days | yes |
| 73 | HCI-057EI | Liver metastasis | Pretreated | pos | pos | pos | pos | pos | pos | No; ESR1 WT | yes | 10–15 days | 25–30 days | yes |
| 74 | HCI-058 | Pleural effusion | Pretreated | neg | neg | neg | neg | neg | neg | n/a | yes | 4–5 days | 15–20 days | yes |
| 75 | HCI-059 | 1° breast | Naïve | neg | neg | neg | neg | neg | neg | n/a | Failed (only mouse content) | | | |
| 76 | HCI-060 | Brain metastasis | Pretreated | neg | neg | neg | neg | neg | neg | n/a | yes | 4–7 days | 15–20 days | yes, also PDO** |
| 77 | HCI-061 | Brain metastasis | Pretreated | neg | neg | neg | neg | neg | neg | n/a | Failed (no organoid formation) | | | |
| 78 | HCI-062 | Residual breast | Pretreated | neg | neg | neg | neg | neg | neg | n/a | yes | 7–10 days | 20–25 days | yes |
| 79 | HCI-063 | Residual breast | Pretreated | neg | neg | neg | neg | neg | neg | n/a | not attempted | | | |
| 80 | HCI-064 | 1° breast | no known tx | pos | pos | pos | pos | pos | pos | Yes; ESR1 WT | yes | 7–10 days | 20–25 days | yes |
| 81 | HCI-065 | Breast recurrence | Naïve | neg | neg | neg | neg | neg | neg | n/a | yes | 7–10 days | 20–25 days | yes |
| 82 | HCI-066 | 1° breast | Naïve | neg | neg | neg | neg | neg | neg | n/a | Failed (no organoid formation) | | | |
| 83 | HCI-067 | 1° breast | Naïve | neg | neg | neg | neg | neg | neg | n/a | yes | 10–15 days | 25–30 days | yes |
| 84 | HCI-068 | Liver metastasis | Pretreated | neg | neg | neg | neg | neg | neg | n/a | yes | 10–12 days | 25–30 days | no |
| 85 | HCI-069 | 1° breast | Naïve | neg | neg | neg | neg | neg | neg | n/a | yes | 7–10 days | 20–25 days | yes |
| 86 | HCI-070 | Brain metastasis | Pretreated | neg | neg | neg | neg | neg | neg | n/a | yes | 10–18 days | 25–30 days | yes |
| 87 | HCI-071 | Brain metastasis | Pretreated | pos (10%) | neg | neg | neg | neg | neg | n/a | yes | 15–20 days | 25–30 days | yes |
| 88 | HCI-072 | Brain metastasis | Pretreated | pos | pos | pos | pos | pos | pos | Yes; ESR1 WT | yes | 4–7 days | 15–20 days | yes |
| 89 | HCI-073 | 1° breast | Naïve | neg | neg | neg | neg | neg | neg | n/a | yes | 10–15 days | 25–30 days | yes |
| 90 | HCI-074 | Residual breast | Pretreated | neg | neg | neg | neg | neg | neg | n/a | yes | 10–15 days | 25–30 days | yes |
| 91 | HCI-075 | Pleural effusion | Pretreated | neg | neg | neg | neg | neg | neg | n/a | yes | 6-10 days | 15-20days | no |
| 92 | HCI-076 | Brain metastasis | Pretreated | neg | neg | pos | neg | neg | pos | n/a | yes | 10–15 days | 25–30 days | yes |
| 93 | HCI-076LN | Brain metastasis | Pretreated | neg | neg | pos | neg | neg | pos | n/a | yes | 10–15 days | 25–30 days | yes |
| 94 | HCI-077 | 1° breast | Naïve | neg | neg | neg | neg | neg | neg | n/a | yes | 7–10 days | 20–25 days | yes |
| 95 | HCI-078 | Residual breast | Pretreated | neg | neg | neg | neg | neg | neg | n/a | yes | 7–10 days | 20–25 days | yes |
| 96 | HCI-079 | Lung metastasis (Phylloides) | Pretreated | neg | neg | neg | neg | neg | neg | n/a | not attempted | | | |
| 97 | HCI-080 | Chest wall | Pretreated | neg | neg | neg | neg | neg | neg | n/a | not attempted | | | |
| 98 | HCI-081 | Brain metastasis | Pretreated | neg | neg | pos | neg | neg | pos | n/a | yes | 8-15 days | 25-30 days | yes |
| 99 | HCI-082 | Brain metastasis | Pretreated | pos | neg | neg | pos | neg | neg | Yes; ESR1 WT | not attempted | | | |
| 100 | HCI-083 | Bone metastasis | Pretreated | neg | neg | neg | pos | neg | neg | Yes; ESR1 WT | not attempted | | | |
| 101 | HCI-084 | 1° breast | Naïve | neg | neg | neg | neg | neg | neg | n/a | not attempted | | | |
| 102 | HCI-085 | Lymph node metastasis | Pretreated | neg | neg | pos | neg | neg | pos | n/a | yes | 7–10 days | 15–20 days | yes |
| 103 | HCI-086 | 1° breast | Naïve | neg | neg | neg | neg | neg | pos | n/a | yes | 7-10 days | 20-25 days | no |
| 104 | HCI-087 | Ascites | Pretreated | pos | pos | neg | pos | pos | neg | No; ESR1 Y537S | yes | 10-12 days | 28-30 days | no |
| 105 | HCI-087EI | Ascites | Pretreated | pos | pos | neg | pos | pos | neg | No; ESR1 Y537S | yes | 11-14 days | 21-25 days | no |
| 106 | HCI-088 | Lung metastasis | Pretreated | neg | neg | neg | neg | neg | neg | n/a | Failed (no organoid formation) | | | |
| 107 | HCI-089 | Brain metastasis | Pretreated | neg | neg | pos | pos | neg | pos | nd; ESR1 WT | yes | 6-10 days | 20-25 days | yes |
| 108 | HCI-090 | Liver metastasis | Pretreated | neg (ER history) | neg | pos | pos | neg | neg | n/a | not attempted | | | |
| 109 | HCI-090EI | Liver metastasis | Pretreated | neg (ER history) | neg | pos | pos | neg | neg | No; ESR1 WT | not attempted | | | |
| 110 | HCI-091 | Brain metastasis | Pretreated | neg | neg | pos | neg | neg | pos | n/a | not attempted | | | |
| 111 | HCI-092 | Lung metastasis | Pretreated | neg | neg | neg | neg | neg | neg | n/a | yes | 5-7 days | 15-25 days | no |
| 112 | HCI-093 | Brain metastasis | Pretreated | neg | neg | neg | neg | neg | neg | n/a | yes | 15-20 days | 30-35 days | no |
| 113 | HCI-094 | Brain metastasis | Pretreated | neg | neg | neg | ongoing | ongoing | ongoing | n/a | not attempted | | | |
| 114 | HCI-095 | Brain metastasis | Pretreated | neg | neg | neg | ongoing | ongoing | ongoing | n/a | not attempted | | | |
| 115 | HCI-096 | Lung metastasis | Pretreated | neg | neg | neg | ongoing | ongoing | ongoing | n/a | not attempted | | | |
| 116 | HCI-097 | Ascites | Pretreated | pos | neg | neg | pos | pos (weak) | neg | ongoing; ESR1 WT | yes | 15-20 days | 25-30 days | no |
| 117 | HCI-097EI | Ascites | Pretreated | pos | neg | neg | ongoing | ongoing | ongoing | ongoing; ESR1 WT | not attempted | | | |
| 118 | HCI-098 | Bone metastasis | Pretreated | neg | neg | neg | neg | neg | neg | n/a | not attempted | | | |
| 119 | HCI-099 | 1° breast | Naïve | neg | neg | neg | neg | neg | neg | n/a | yes | 10-15 days | 25-30 days | no |
| 120 | HCI-100 | Pleural effusion | Pretreated | pos | pos | neg | pos | pos | neg | ongoing | not attempted | | | |
| 121 | HCI-101 | Brain metastasis (Phylloides) | Pretreated | neg | neg | neg | neg | neg | neg | n/a | not attempted | | | |
| 122 | HCI-102 | Liver metastasis | Pretreated | pos | pos | pos | ongoing | ongoing | pos | nd; ESR1 WT | not attempted | | | |
| 123 | HCI-103 | Brain metastasis | Pretreated | neg | neg | pos | neg | neg | pos | n/a | not attempted | | | |
| 124 | HCI-104 | Brain metastasis | Pretreated | pos | pos | neg | pos | pos | neg | ongoing | not attempted | | | |
| 125 | HCI-105 | 1° breast | Naïve | neg | neg | neg | neg | neg | neg | n/a | not attempted | | | |
| 126 | HCI-106 | LN metastasis | Pretreated | neg | neg | neg | neg | neg | neg | n/a | not attempted | | | |
| 127 | TOW-85 | 1° breast | Naïve | neg | neg | neg | ongoing PDXOX | ongoing PDXOX | ongoing PDXOX | n/a | yes | 7-10 days | 20-25 days | yes |
| 128 | BCM-0132 | 1° breast | Naïve | neg | neg | neg | neg | neg | neg | n/a | yes | 7-10 days | 12-15 days | yes |
| 129 | BCM-2665 | Breast | Pretreated | neg | neg | neg | neg | neg | neg | n/a | yes | 7-10 days | 12-15 days | yes |
| 130 | BCM-3143 | Breast | Pretreated | neg | neg | pos | neg | neg | pos | n/a | yes | 7-10 days | 12-15 days | yes |
| 131 | BCM-3277 | Breast | Pretreated | pos | pos | neg | pos | pos | neg | Yes; ESR1 WT | yes | 7-10 days | 12-15 days | yes |
| 132 | BCM-3561 | Ascites | Pretreated | pos | pos | neg | neg | neg | neg | n/a | yes | 4-7 days | 12-15 days | yes |
| 133 | BCM-3611 | 1° breast | Naïve | neg | neg | neg | neg | neg | neg | n/a | yes | 4-7 days | 12-15 days | yes |
| 134 | BCM-3904 | Breast | Naïve | neg | neg | neg | neg | neg | neg | n/a | yes | 4-7- days | 12-15 days | yes |
| 135 | BCM-3936 | Breast | Pretreated | neg | neg | neg | neg | neg | neg | n/a | yes | 10-15 days | 20-25 days | yes |
| 136 | BCM-3963 | 1° breast | Naïve | neg | neg | pos | neg | neg | pos | n/a | yes | 7-10 days | 12-15 days | yes |
| 137 | BCM-4175 | Breast | Pretreated | neg | neg | neg | neg | neg | neg | n/a | yes | 4-7 days | 12-15 days | yes |
| 138 | BCM-4195 | Breast | Pretreated | neg | neg | neg | neg | neg | neg | n/a | yes | 10-15 days | 20-25 days | yes |
| 139 | BCM-4272 | 1° breast | Naïve | neg | neg | neg | neg | neg | neg | n/a | yes | 7-10 days | 15-20 days | yes |
| 140 | BCM-4888 | Breast | Pretreated | pos | pos | neg | pos | pos | pos | Yes; ESR1 WT | yes | 10-15 days | 25-30 days | yes |
| 141 | BCM-4913 | Breast | Pretreated | neg | neg | neg | neg | neg | neg | n/a | yes | 4-7 days | 12-15 days | yes |
| 142 | BCM-5471 | Breast | Pretreated | unknown | unknown | unknown | neg | neg | neg | n/a | yes | 4-7 days | 12-15 days | yes |
| 143 | BCM-5998 | Chest wall | Pretreated | neg | neg | neg | neg | neg | neg | n/a | yes | 4-7 days | 15-20 days | yes |
| 144 | BCM-7441 | Breast | Pretreated | pos | pos | neg | pos | neg | neg | Yes; ESR1 WT | yes | 4-7 days | 20-25 days | yes |
| 145 | BCM-7482 | Breast | Pretreated | neg | neg | neg | neg | neg | neg | n/a | yes | 10-15 days | 20-25 days | yes |
| 146 | BCM-7821 | Breast | Pretreated | neg | neg | neg | neg | neg | neg | n/a | yes | 7-10 days | 15-20 days | yes |
| 147 | BCM-15006 | Skin metastasis | Pretreated | pos | neg | neg | neg | neg | neg | n/a | yes | 7-10 days | 15-20 days | no |
| 148 | BCM-15034 | Liver metastasis | Pretreated | pos | neg | neg | pos | pos | unknown | Yes; ESR1 WT | yes | 7-10 days | 15-20 days | yes |
| 149 | BCM-15057 | Breast | Pretreated | pos | neg | neg | pos | pos | neg | Yes; ESR1 WT | yes | 10-15 days | 20-25 days | yes |
| 150 | BCM-15100 | Liver metastasis | Pretreated | pos | pos | neg | pos | pos | neg | Yes; ESR1 WT | yes, but grew too slow | | | |
| 151 | BCM-15133 | Bone metastasis | Pretreated | pos | neg | neg | pos | pos | neg | Yes; ESR1 WT | yes | 10-15 days | 20-25 days | yes |
| 152 | BCM-15156 | Breast | Pretreated | pos | neg | neg | pos | neg | neg | Yes; ESR1 WT | yes | 7-10 days | 20-25 days | yes |
| 153 | BCM-15160 | Pleural effusion | Pretreated | pos | neg | neg | pos | neg | neg | Yes; ESR1 WT | yes | 7-10 days | 20-25 days | yes |
| 154 | BCM-15163 | Breast | Naïve | neg | neg | pos | neg | neg | pos | n/a | yes | 7-10 days | 20-25 days | yes |
| 155 | BCM-15171 | Lymph node metastasis | Pretreated | neg | neg | neg | neg | neg | pos | n/a | yes | 7-10 days | 20-25 days | yes |
| 156 | BCM-15214 | Breast | Pretreated | neg | neg | neg | pos | pos | pos | Yes; ESR1 WT | yes | 10-15 days | 20-25 days | yes |

Abbreviations: 1°, primary; pos, positive; neg, negative; n/a, not applicable; nd, not done; E2, estradiol; PDX, Patient-derived xenograft; PDXO, PDX-derived organoids; recurr., recurrent; LN, lymph node; BRN, Brain met; LV, Liver met; OV, ovary met; tx, treatment; FR, fulvestrant resistant; BR, birinapant resistant; BS, birinapant sensitive; BGR, birinapant gemcitabine resistant; VR, vinorelbine resistant; CR, capecitabine resistant

Model IDs with matching color models indicate multiple models from the same patient

* PDXOX (PDX established from injeciton of PDXOs)

** PDO (patient-derived organoid) was develped in Clevers media following Sachs et al., 2018
