## Supplementary Table S2. Summary of raw PDXO screening data for "A precision oncology-focused deep learning framework for personalized selection of cancer therapy"

Drug

PDxO Model

| | ERK MAPK signaling | | |
|---|---|---|---|
| Drug | Dabrafenib | Selumetinib | Sotorasib |
| Target | B-raf | MEK1/2 | KRAS G12C |
| Screen | A | ABCD | C |
| HCI-001 | X | X | |
| HCI-002 | X | X | |
| HCI-003 | X | X | |
| HCI-005 | X | X | |
| HCI-007 | X | X | |
| HCI-008 | | X | X |
| HCI-010 | X | X | |
| HCI-011 | X | X | |
| HCI-012 | X | X | |
| HCI-015 | X | X | |
| HCI-015BGR | | X | |
| HCI-015BR | | X | |
| HCI-016 | X | X | |
| HCI-017 | X | X | |
| HCI-018 | | X | X |
| HCI-019 | X | X | |
| HCI-023 | X | X | |
| HCI-023BR | | X | |
| HCI-024 | | X | X |
| HCI-025 | X | X | |
| HCI-027 | X | X | |
| HCI-027BR | X | X | |
| HCI-027BS | | X | |
| HCI-028 | | X | X |
| HCI-028LV | | X | |
| HCI-031 | | X | |
| HCI-032 | X | X | |
| HCI-032EI | | X | X |
| HCI-037 | | X | |
| HCI-038 | | X | |
| HCI-039 | | X | |
| HCI-040 | | X | |
| HCI-040EI | | X | |
| HCI-041 | | X | |
| HCI-042 | | X | |
| HCI-043 | X | X | |
| HCI-045 | X | X | |
| HCI-045VR | | X | |
| HCI-046 | | X | |
| HCI-047A2 | | X | X |
| HCI-048 | | X | |
| HCI-048CR | | X | |
| HCI-050 | | X | X |
| HCI-051 | | X | |
| HCI-053 | | X | X |
| HCI-053CR | | X | |
| HCI-054 | | X | |
| HCI-054CR | | X | |
| HCI-055 | | X | X |
| HCI-056 | | X | |
| HCI-057 | | X | |
| HCI-057EI | | X | X |
| HCI-058 | | X | |
| HCI-060 | | X | |
| HCI-060PDO | | X | |
| HCI-062 | | X | X |
| HCI-064 | | X | |
| HCI-065 | | X | X |
| HCI-067 | | X | X |
| HCI-069 | | X | |
| HCI-070 | | X | |
| HCI-071 | | X | X |
| HCI-072 | | X | X |
| HCI-073 | | X | X |
| HCI-074 | | X | |
| HCI-076 | | X | X |
| HCI-076LN | | X | |
| HCI-077 | | X | X |
| HCI-078 | | X | X |
| HCI-081 | | X | |
| HCI-085 | | X | |
| HCI-089 | | X | |
| TOW-85 | | X | |
| BCM-0132 | | X | X |
| BCM-2665 | | X | |
| BCM-3143 | X | X | |
| BCM-3277 | | X | |
| BCM-3561 | | X | X |
| BCM-3611 | | X | |
| BCM-3904 | X | X | |
| BCM-3936 | | X | |
| BCM-3963 | X | X | |
| BCM-4175 | | X | |
| BCM-4195 | | X | |
| BCM-4272 | | X | |
| BCM-4888 | X | X | |
| BCM-4913 | | X | X |
| BCM-5471 | X | X | |
| BCM-5998 | | X | |
| BCM-7441 | | X | |
| BCM-7482 | | X | |
| BCM-7821 | X | X | |
| BCM-15034 | | X | X |
| BCM-15057 | | X | X |
| BCM-15133 | | X | |
| BCM-15156 | | X | |
| BCM-15160 | | X | X |
| BCM-15163 | | X | |
| BCM-15171 | | X | |
| BCM-15214 | | X | |

| | ERK MAPK signaling | DNA replication | | | Cell cycle | | | | Chromatin histone methylation | Genome integrity | | | | | | | | | | | Mitosis | | | | Metabolism | Apoptosis regulation | | | | | | | | | | | | EGFR signaling | | | | Protein stability and degradation | | | | Hormone-related | | | | Chromatin histone acetylation | | | WNT signaling | p53 pathway | PI3K/MTOR signaling | | | | | | | | RTK signaling | | | | | | | | Other, kinases | | | | | | | | | | | |
|---|---|---|---|---|---|---|---|---|---|---|---|---|---|---|---|---|---|---|---|---|---|---|---|---|---|---|---|---|---|---|---|---|---|---|---|---|---|---|---|---|---|---|---|---|---|---|---|---|---|---|---|---|---|---|---|---|---|---|---|---|---|---|---|---|---|---|---|---|---|---|---|---|---|---|---|---|---|---|---|---|---|---|
| Drug | Trametinib | Carboplatin | Methoxyamine | Triapine | Abemaciclib | AZD4573 | MK-1775 | Palbociclib | EPZ5676 | A-966492 | AZD0156 | Iniparib | Nedisertib | Niraparib | Olaparib | Pamiparib | Pevonedistat | Rucaparib | Talazoparib | VE-822 | Alisertib | Docetaxel | Eribulin | Vinorelbine | Epacadostat | APG-1387 | ASTX660 | AZD5991 | Birinapant | GDC-0152 | GDC-0917 | LCL161 | MIK665 | Navitoclax | Pelcitoclax | Tucatinib | Venetoclax | Afatinib | Lapatinib | Neratinib | Osimertinib | Bortezomib | Ixazomib | Onalespib | TAK-243 | Endoxifen | Enzalutamide | Fulvestrant | Megestrol Acetate | Belinostat | Entinostat | Romidepsin | RO4929097 | AMG232 | Alpelisib | AZD5363 | Copanlisib | Everolimus | Ipatasertib | MK-2206 | Sapanisertib | Taselisib | AMG337 | Cabozantinib | Crizotinib | Erdafitinib | PD173074 | Savolitinib | Sunitinib | Tivantinib | 5-Aza | 4μ8C | Dasatinib | Emavusertib | EPZ011989 | Ibrutinib | LDC204857 | ONC206 | Ruxolitinib | Selinexor | Sorafenib | Telaglenastat |
| Target | MEK | DNA synthesis | DNA synthesis | RNR | CDK4/6 | CDK9 | Wee1 | CDK4/6 | DOT1L | PARP1/2 | ATM | PARP1 | DNA-PK | PARP1/2 | PARP1/2 | PARP1/2 | NAE | PARP1/2/3 | PARP1/2 | ATR/ATM | Aurora A | Microtubule | Microtubule | Microtubule | IDO1 | IAP | cIAP,XIAP | Mcl-1 | cIAP,XIAP | IAP | IAP | IAP | Mcl-1 | Bcl-2 | Bcl-2/Bcl-xl | HER2 | Bcl-2 | EGFR | HER2,EGFR | HER2,EGFR | EGFR | Proteasome | Proteasome | Hsp90 | UAE | ER | AR | ER | PR | HDAC | HDAC | HDAC | γ secretase | MDM2 | PI3Kα | Akt | PI3K | mTOR1 | Akt | Akt | mTOR1/2 | PI3K | MET | VEGFR2,MET | c-Met,ALK | FGFR | FGFR1 | c-Met | VEGFR2,PDGFRβ | c-Met | DNMT1 | IRE1 | Src/Bcr-Abl | IRAK4/FLT3 | EZH2 | Btk | POLRMT | DRD2/3/4 | JAK1/2 | CRM1 | Raf | GLS1 |
| Screen | BCD | ABCD | A | A | ABCD | CD | AB | A | A | B | A | B | AB | A | ABCD | BCD | ABCD | BCD | ABCD | ABCD | BCD | ABCD | ABCD | ABCD | C | D | BCD | CD | ABC | B | B | B | CD | ABCD | D | B | A | BCD | AB | ABCD | C | A | ABCD | BCD | AB | CD | A | A | A | CD | AB | A | A | AB | BCD | D | ABCD | ABCD | B | ABCD | BCD | A | A | A | A | BCD | C | A | C | ABCD | BCD | A | A | C | A | CD | D | D | AB | CD | A | A |
| HCI-001 | | X | X | X | X | | X | X | X | | X | | X | X | X | | X | | X | X | | X | X | X | | | | | X | | | | | X | | | X | | X | X | | X | X | | X | | X | X | X | | X | X | X | X | | | X | X | | X | | X | X | X | X | | | X | | X | | | X | | X | | | | X | | X | X |
| HCI-002 | | X | X | X | X | | X | X | X | | X | | X | X | X | | X | | | X | | X | | | | | | | X | | | | | X | | | X | | X | X | | X | X | | X | | | X | X | | X | X | X | X | | | X | X | | | | X | X | X | X | | | X | | X | | X | X | | X | | | | X | | X | X |
| HCI-003 | | X | X | X | X | | X | X | X | | X | | X | X | X | | X | | X | X | | X | X | X | | | | | X | | | | | X | | | X | | X | X | | X | X | | X | | X | X | X | | X | X | X | X | | | X | X | | X | | X | X | X | X | | | X | | X | | X | X | | X | | | | X | | X | X |
| HCI-005 | | X | X | X | X | | X | X | X | | X | | X | X | X | | X | | X | X | | X | X | X | | | | | X | | | | | X | | | X | | X | X | | X | X | | X | | X | X | X | | X | X | X | X | | | X | X | | X | | X | X | X | X | | | X | | X | | X | X | | X | | | | X | | X | X |
| HCI-007 | | X | X | X | X | | X | X | X | | X | | X | X | X | | X | | X | X | | X | X | X | | | | | X | | | | | X | | | X | | X | X | | X | X | | X | | X | X | | | X | X | X | X | | | X | X | | X | | X | X | X | X | | | X | | X | | X | X | | X | | | | X | | X | X |
| HCI-008 | X | X | | | X | X | | | | | | | | | X | X | X | X | X | X | X | X | X | X | X | | X | X | X | | | | X | X | | | | X | | X | X | | X | X | | X | | | | X | | | | | X | | X | X | | X | X | | | | | X | X | | X | X | X | | | X | | X | | | | X | | |
| HCI-010 | [illegible] | [illegible] | [illegible] | [illegible] | [illegible] | [illegible] | [illegible] | [illegible] | [illegible] | [illegible] | [illegible] | [illegible] | [illegible] | [illegible] | [illegible] | [illegible] | [illegible] | [illegible] | [illegible] | [illegible] | [illegible] | [illegible] | [illegible] | [illegible] | [illegible] | [illegible] | [illegible] | [illegible] | [illegible] | [illegible] | [illegible] | [illegible] | [illegible] | [illegible] | [illegible] | [illegible] | [illegible] | [illegible] | [illegible] | [illegible] | [illegible] | [illegible] | [illegible] | [illegible] | [illegible] | [illegible] | [illegible] | [illegible] | [illegible] | [illegible] | [illegible] | [illegible] | [illegible] | [illegible] | [illegible] | [illegible] | [illegible] | [illegible] | [illegible] | [illegible] | [illegible] | [illegible] | [illegible] | [illegible] | [illegible] | [illegible] | [illegible] | [illegible] | [illegible] | [illegible] | [illegible] | [illegible] | [illegible] | [illegible] | [illegible] | [illegible] | [illegible] | [illegible] | [illegible] | [illegible] | [illegible] | [illegible] |
| HCI-011 | [illegible] | [illegible] | [illegible] | [illegible] | [illegible] | [illegible] | [illegible] | [illegible] | [illegible] | [illegible] | [illegible] | [illegible] | [illegible] | [illegible] | [illegible] | [illegible] | [illegible] | [illegible] | [illegible] | [illegible] | [illegible] | [illegible] | [illegible] | [illegible] | [illegible] | [illegible] | [illegible] | [illegible] | [illegible] | [illegible] | [illegible] | [illegible] | [illegible] | [illegible] | [illegible] | [illegible] | [illegible] | [illegible] | [illegible] | [illegible] | [illegible] | [illegible] | [illegible] | [illegible] | [illegible] | [illegible] | [illegible] | [illegible] | [illegible] | [illegible] | [illegible] | [illegible] | [illegible] | [illegible] | [illegible] | [illegible] | [illegible] | [illegible] | [illegible] | [illegible] | [illegible] | [illegible] | [illegible] | [illegible] | [illegible] | [illegible] | [illegible] | [illegible] | [illegible] | [illegible] | [illegible] | [illegible] | [illegible] | [illegible] | [illegible] | [illegible] | [illegible] | [illegible] | [illegible] | [illegible] | [illegible] | [illegible] |
| HCI-012 | [illegible] | [illegible] | [illegible] | [illegible] | [illegible] | [illegible] | [illegible] | [illegible] | [illegible] | [illegible] | [illegible] | [illegible] | [illegible] | [illegible] | [illegible] | [illegible] | [illegible] | [illegible] | [illegible] | [illegible] | [illegible] | [illegible] | [illegible] | [illegible] | [illegible] | [illegible] | [illegible] | [illegible] | [illegible] | [illegible] | [illegible] | [illegible] | [illegible] | [illegible] | [illegible] | [illegible] | [illegible] | [illegible] | [illegible] | [illegible] | [illegible] | [illegible] | [illegible] | [illegible] | [illegible] | [illegible] | [illegible] | [illegible] | [illegible] | [illegible] | [illegible] | [illegible] | [illegible] | [illegible] | [illegible] | [illegible] | [illegible] | [illegible] | [illegible] | [illegible] | [illegible] | [illegible] | [illegible] | [illegible] | [illegible] | [illegible] | [illegible] | [illegible] | [illegible] | [illegible] | [illegible] | [illegible] | [illegible] | [illegible] | [illegible] | [illegible] | [illegible] | [illegible] | [illegible] | [illegible] | [illegible] | [illegible] |
| HCI-015 to BCM-15214 (remaining 91 rows) | [illegible] | [illegible] | [illegible] | [illegible] | [illegible] | [illegible] | [illegible] | [illegible] | [illegible] | [illegible] | [illegible] | [illegible] | [illegible] | [illegible] | [illegible] | [illegible] | [illegible] | [illegible] | [illegible] | [illegible] | [illegible] | [illegible] | [illegible] | [illegible] | [illegible] | [illegible] | [illegible] | [illegible] | [illegible] | [illegible] | [illegible] | [illegible] | [illegible] | [illegible] | [illegible] | [illegible] | [illegible] | [illegible] | [illegible] | [illegible] | [illegible] | [illegible] | [illegible] | [illegible] | [illegible] | [illegible] | [illegible] | [illegible] | [illegible] | [illegible] | [illegible] | [illegible] | [illegible] | [illegible] | [illegible] | [illegible] | [illegible] | [illegible] | [illegible] | [illegible] | [illegible] | [illegible] | [illegible] | [illegible] | [illegible] | [illegible] | [illegible] | [illegible] | [illegible] | [illegible] | [illegible] | [illegible] | [illegible] | [illegible] | [illegible] | [illegible] | [illegible] | [illegible] | [illegible] | [illegible] | [illegible] | [illegible] |
